# Systematic genome-wide querying of coding and non-coding functional elements in *E. coli* using CRISPRi

**DOI:** 10.1101/2020.03.04.975888

**Authors:** Harneet S. Rishi, Esteban Toro, Honglei Liu, Xiaowo Wang, Lei S. Qi, Adam P. Arkin

**Affiliations:** Biophysics Graduate Group, University of California - Berkeley, Berkeley, CA, 94720, USA; Designated Emphasis Program in Computational and Genomic Biology, University of California - Berkeley, Berkeley, CA, 94720, USA; Department of Bioengineering, University of California - Berkeley, Berkeley, CA, 94720, USA; Bioinformatics Division, Center for Synthetic and Systems Biology, Tsinghua National Laboratory for Information Science and Technology; Department of Automation, Tsinghua University; Department of Bioengineering, Stanford University, Stanford, CA, 94305, USA; Department of Chemical and Systems Biology, Stanford University, Stanford, CA, 94305, USA; Stanford ChEM-H, Stanford University, Stanford, CA 94305, USA; Environmental Genomics and Systems Biology Division, Lawrence Berkeley National Laboratory, Berkeley, CA, 94720, USA

## Abstract

Genome-wide repression screens using CRISPR interference (CRISPRi) have enabled the high-throughput identification of essential genes in bacteria. However, there is a lack of functional studies leveraging CRISPRi to systematically explore targeting of both the coding and non-coding genome in bacteria. Here we perform CRISPRi screens in *Escherichia coli* MG1655 K-12 targeting ~13,000 genomic features, including nearly all protein-coding genes, non-coding RNAs, promoters, and transcription factor binding sites (TFBSs) using a ~33,000-member sgRNA library, which represents the most compact and comprehensive genome-wide CRISPRi library in *E. coli* to date. Our data reveal insights into the conditional essentiality of the genome with key refinements to screen design and profiling. First, we demonstrate that strong fitness defects associated with essential cellular processes can be resolved using inducible time-series measurements. We show that knockdowns of different classes of genes exhibit distinct, transient responses that are correlated to gene function with genes involved in translation exhibiting the strongest responses. We also query feature essentiality across several biochemical conditions and show that several genes, sRNAs, and operons exhibit conditional phenotypes not reported by previous high-throughput efforts. Second, we evaluate systematically targeting non-genic features (promoters and TFBSs) in the *E. coli* genome. We show that promoter-targeting guides can be used to add phenotypic confidence to promoter annotations and verify computationally predicted promoters. In contrast to prior studies, we find that promoter knockdowns exhibit a strong targeting orientation dependency where targeting the non-template strand of the promoter closest to the target gene is more effective in knocking down gene expression than other promoter targeting orientations. Unlike eukaryotic genomes, we note that interpreting the effects of TFBS targeting is particularly challenging due to the small size of such features and their proximity to and overlap with other genomic features. Together, this work reveals novel conditionally essential gene phenotypes, provides a characterized set of sgRNAs for future *E. coli* CRISPRi screens, and highlights considerations for CRISPRi library design and screening for microbial genome characterization.

## INTRODUCTION

The nuclease deactivated dCas9 protein has been developed as a powerful tool for programmable gene repression^1^, and the ability to induce genetic perturbation at a user-defined time – a feature not available in conventional gene disruption or deletion techniques – has enabled the CRISPRi-mediated characterization of essential genes in a number of bacteria^2-7^. The programmability of CRISPRi targeting also enables the interrogation of smaller non-coding DNA (ncDNA) features such as non-coding RNA (ncRNA) genes, promoters, and transcription factor binding sites (TFBSs). ncDNA features, which represent ~12 percent of the *E. coli* genome, play important roles in the regulation of gene expression in a condition-dependent manner. For example, small RNAs (sRNAs) have been implicated in transient regulatory processes involving membrane biogenesis, metabolism, and the synthesis of key transcription factors^8^ while ncDNA regulatory elements drive key physiological decisions such as complex metabolism^9^, pathogenicity^10^, and gene expression diversification^11^. However, ncDNA features have been difficult to perturb using traditional genome-scale methods (e.g. targeted modifications using λ-Red recombination^12-16^, random insertions using transposon elements^17-20^) due to the random targeting of transposons and disruption of local genomic context by insertions, making their interrogation via CRISPRi highly valuable.

Despite this potential value-add, previous bacterial CRISPRi screening studies have been limited in their investigation of RNA genes beyond simple cases (e.g. tRNA, rRNA genes) and have rarely addressed non-coding genomic features such as promoters and TFBSs. In comparison, CRISPRi screens in eukaryotic systems have been routinely employed to find new regulatory sites in enhancer regions^21-23^ and functionally profile lncRNAs^24-27^, indicating the untapped potential of CRISPRi for the functional characterization of bacterial genomes. In addition, existing CRISPRi screens measure phenotypes using end-point fitness measurements by calculating the change in strain abundance between the beginning and end of a screen, which ignores dynamic outcomes that may occur over the course of an experiment. However, the physiological response resulting from CRISPRi-mediated gene repression could vary between different genes, arising from differences in protein and mRNA decay rates, feedback regulation, interaction network structure, and the physiological relevance of the targeted gene itself. Resolving such end-point measurements by tracking transient responses upon CRISPRi knockdown can yield rich insights into the primacy of processes for fitness and highlight the time it takes to demonstrate a physiological effect in different environments.

Here we leverage the programmable nature of CRISPRi to target approximately 13,000 *E. coli* MG1655 K-12 genomic features (protein-coding genes, non-coding RNAs (ncRNAs), promoters, and TFBSs) using a compact, designed oligo array library of 32,992 sgRNAs. We first validated our technology by showing that we could knock down 90% of essential genes (as annotated by the Profiling of *E. coli* Chromosome - PEC - database^28,29^) in a pooled screen with the entire library. Through this process, we showed that a designed, compact library with ~4 guides/gene is sufficient for probing gene essentiality, which represents a considerable reduction in comparison to a previous designed *E. coli* screening study using 15 guides/gene^4^. Given that gene essentiality is context-dependent, we expected that querying essentiality under a variety of biochemical conditions would allow us to delineate between a core set of essential genes and an accessory set of conditionally-essential genes. We thus leveraged the inducible nature of CRISPRi to propagate strains targeting essential genomic features and assay the library in several conditions to find condition-dependent phenotypes for essential genes and ncRNAs. Having validated the library, we next investigated if endpoint phenotypes could be further resolved by investigating how different genomic features respond upon CRISPRi induction. We used time-series measurements to track the dynamic response of genes in our library to CRISPRi perturbation and showed that essential genes exhibited distinct profiles that were correlated with their physiological function – a phenomenon not reported from previous CRISPRi screens due to their use of only endpoint measurements of fitness.

Finally, we studied the physiological effects of perturbing DNA regulatory elements such as promoters and TFBSs as these features have been understudied in previous bacterial CRISPRi screens. We showed that targeting promoters of essential genes could knock down gene expression and used this phenotypic outcome to add annotation strength to RegulonDB promoters. We also showed that perturbing gene expression was more successful when inhibiting transcription elongation (gene targeting CRISPRi) as opposed to inhibiting transcription initiation (promoter targeting CRISPRi) in our library through a comparison of guides targeting the promoter and gene sequences of known essential genes. By analyzing differences in sgRNA design features and the genomic context of targeted promoters, we found targeting the non-template strand of the promoter closest to the target gene was more effective in knocking down gene expression than other promoter targeting orientations, indicating a new design consideration for promoter CRISPRi. Finally, we looked at the effect of dCas9 targeting to TFBSs to see if TFBS-targeted CRISPRi could perturb gene expression. We analyzed TFBSs regulating promoters of essential genes; however, due to the proximity or overlap of targeted TFBSs with promoters we were unable to associate phenotypes to specific TFBS features in most cases – finding only one case of a condition-dependent phenotype for a TFBS cluster regulating the expression of a conditionally-essential aerobic respiration gene. Together, this work represents an extension and characterization of bacterial CRISPRi screens as well as a framework for the design, construction, pooled screening, and analysis of CRISPRi libraries for the high-throughput functional annotation of bacterial genomic features.

## RESULTS

### Design and Construction of CRISPRi Library

We designed a CRISPRi library consisting of 32,992 unique sgRNAs to target 4,457 genes (including 130 small RNAs; sRNAs) and gene-like elements (e.g. insertion elements / prophages), 7,442 promoters and transcription start sites (TSSs), and 1,060 transcription factor binding sites (TFBS) across the *E. coli* K-12 MG1655 genome using bioinformatic and biophysical design constraints (Fig. 1a, see Supplementary Note 1 for design details, Supplementary Table 1a for sequences). In brief, guides were designed to target proximal to a protospacer adjacent motif (PAM) site (NGG for *S. pyogenes* dCas9 used in this work), target a unique genomic sequence, maintain the secondary structure of the sgRNA, and avoid extreme GC content (GC < 20%, GC > 80%), where possible. Gene-targeting guides were designed to target the nontemplate strand and target close to the beginning of the gene, following previous observations ^1,30^. When possible, multiple guides were designed for each feature. The designed sgRNAs were synthesized as an oligo pool (Supplementary Table 1b). To allow for the screening of smaller, more focused libraries the terminal 3’ end of each oligo was designed with a category code that allows for the amplification of subsets from the oligo library (Fig. 1b, Supplementary Table 1c). To construct the genome-wide library, sgRNAs were PCR amplified from the oligo pool and then cloned into an expression vector using a golden gate assembly strategy (Methods). This expression vector (ColE1 origin) maintains the guides under arabinose-inducible control using a pBad promoter. The sgRNA library assembly was transformed into a strain harboring a genomically-encoded *dCas9* under aTc-inducible control using a pTet promoter.

**Fig. 1.**
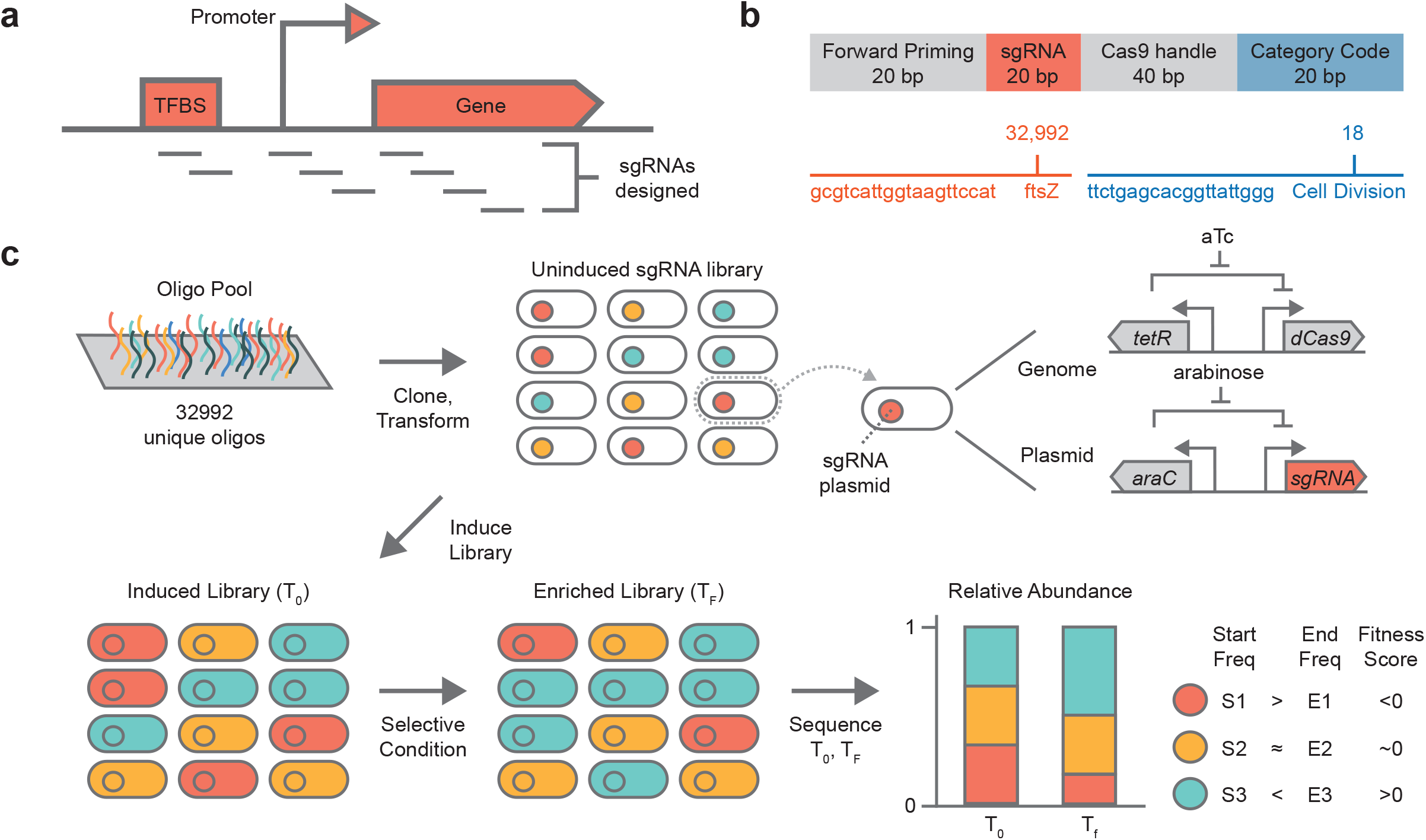
Overview of CRISPRi screening platform. **a** Guide sequences were designed to target three feature types on the *E. coli* genome: (i) gene sequences (ii) promoters (iii) transcription factor binding sites (TFBSs). Multiple guides were designed for each feature where possible (Methods). **b** Guide sequences were synthesized as oligos and ordered via Agilent Technologies as a pool. Category codes (short DNA barcodes) were included in designed oligos to enable amplification of subpools from the library. **c** Guides were first cloned into a receiver vector and transformed into a strain containing chromosomally integrated *dCas9*. At the beginning of an experiment the library is induced and an initial time-point (T_0_) is taken. After growth in a selective condition for a period of time a final time-point (T_F_) is taken. The initial and final samples are sequenced and the fitness of each library member is calculated.

The identity of each knockdown strain in the library is determined solely by the sgRNA plasmid it harbors, specifically the 20 base pair variable region of the sgRNA that directs dCas9 targeting and encodes a DNA barcode for the strain. The relative abundance of every sgRNA, and by extension every strain, can be measured by amplicon sequencing of the variable sgRNA region from a plasmid DNA extraction of the sgRNA library. To perform a pooled functional screen the library is induced to express dCas9 and the sgRNAs and grown under selection for a short period of time (e.g. 24 population doublings) in a user-defined experimental condition (Fig. 1c, Methods). During this competition, strains that carry an sgRNA targeting a feature important for growth will decrease in abundance in the pool. This phenotypic outcome can be quantified by measuring the starting and ending frequency of each strain and calculating a fitness score, which is defined as the normalized log2 ratio of the relative abundance of the guide-strain after the experiment to before the experiment (Methods). For gene targeting guides, we also define a composite gene fitness score as the median of fitness scores for all guides targeting a gene.

### Technology Validation of Genome-Wide CRISPRi Gene Knockdowns

To assess the ability of the library to yield biologically meaningful results, we profiled the phenotypic effect of knockdown for all genes in the library via a fitness experiment in LB Lennox rich media (LB). We found that CRISPRi was highly reproducible (Pearson r_bioiogical_ = 0.90, p < 0.05, permutation test; Pearson r_technical_ = 0.96, p < 0.05, permutation test) (Supplementary Fig. 1). Furthermore, we observed that sgRNAs targeting known essential genes were severely depleted (i.e. strains harboring these guides exhibited a strong growth defect) over the course of an experiment when compared with sgRNAs targeting non-essential genes (Fig. 2a). We compared the fitness results with the Profiling of E. coli Chromosome (PEC) database, which reports 304 E. coli K-12 MG1655 genes for which a knockout could not be generated, implying that these genes were essential for growth in LB rich medium under aerobic conditions (i.e. the condition of library construction) ^28,29^. sgRNAs targeting 274 of 303 (~90%) essential genes were severely depleted (composite gene fitness ≤ -2) over the course of CRISPRi fitness experiments in the same condition, yielding 90 percent agreement with the PEC database. This also included prope
r depletion of all essential E. coli ncRNAs assayed in the experiment as well. Of the remaining 29 essential genes, 15 had at least one sgRNA with fitness ≤ -2 and an additional six had at least one sgRNA with fitness ≤ -1 (Supplementary Table 2). Overall, we found that 289 of 303 essential genes (~95%) could be knocked down by at least one designed sgRNA with fitness ≤ -2, indicating high activity of the CRISPRi library. We also tested the library in M9 minimal medium (M9) under aerobic conditions and found that 385 out of 415 (93%) minimal media essential genes had a gene fitness score ≤ -2 when knocked down (Supplementary Fig. 2).

**Fig. 2.**
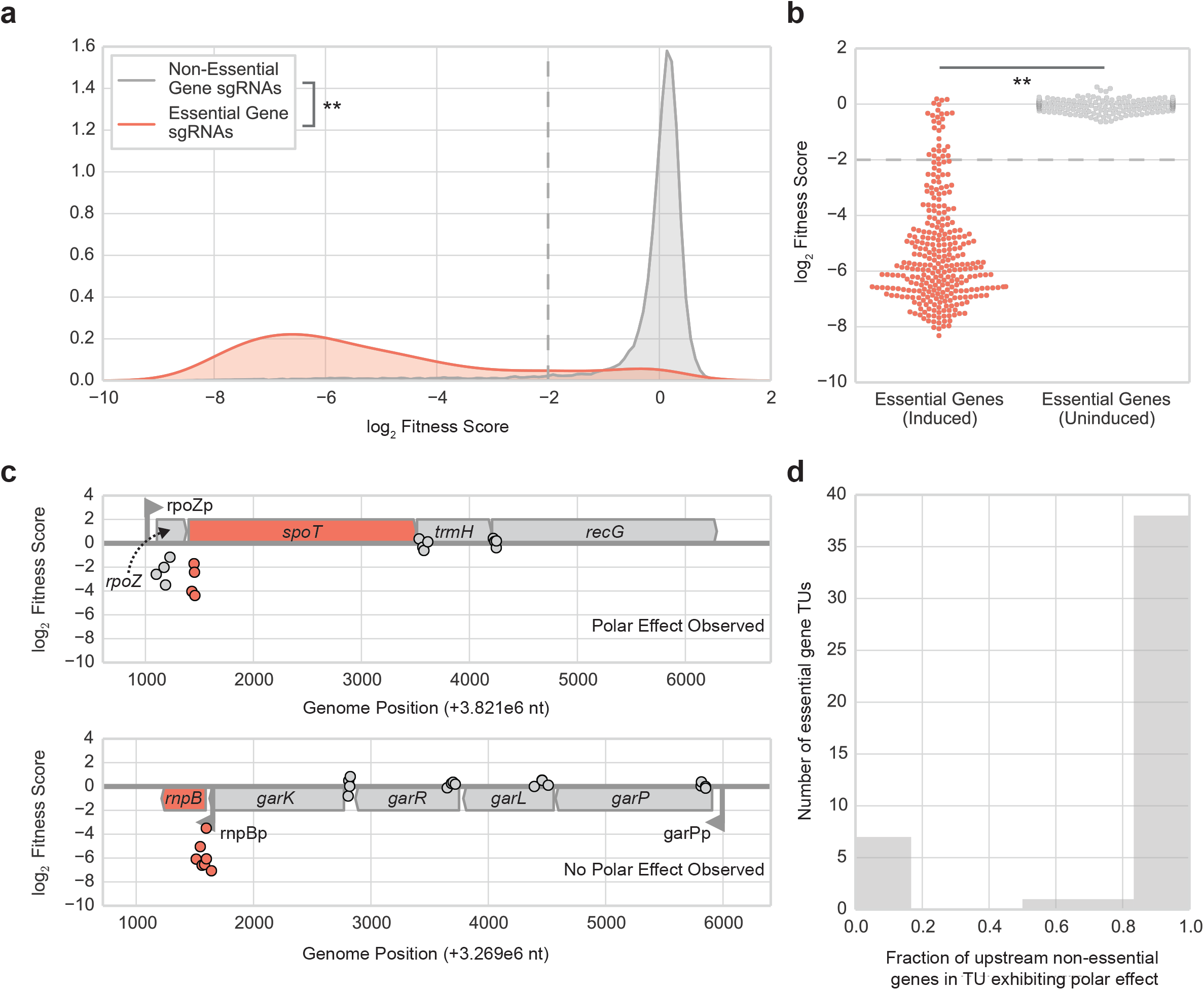
Technology validation of CRISPRi screening platform. **a** Depletion of essential gene targeting sgRNAs compared to non-essential gene targeting sgRNAs over the course of a pooled fitness experiment with the CRISPRi library in LB rich media (with CRISPRi system induced) under aerobic growth conditions for 24 population doublings. **p < 0.001 (Mann-Whitney U-test); Cohen’s *d* = 3.7. **b** Demonstration of tight, inducible control of sgRNA library via comparison of essential gene fitness scores from pooled fitness experiments where the CRISPRi library was either induced (left) or uninduced (right). In the induced condition, the library was induced with aTc and arabinose to express *dCas9* and sgRNA and then grown in LB media for 24 doublings as in a regular fitness experiment (Methods). In the uninduced condition, the library was also grown in similar culturing conditions (e.g. LB media for 24 doublings); however, neither aTc nor arabinose were added. **p < 0.001 (Mann-Whitney U-test); Cohen’s *d* = 3.7. **c** Example of CRISPRi-mediated polar operon effects where targeting a non-essential gene (*rpoZ*) upstream of an essential gene (*spoT*) in the same transcriptional unit (*rpoZ-spoT-trmH-recG*) produces a fitness defect (top panel). In the presence of an intra-operonic promoter (eg rnpBp), knockdown of upstream non-essential genes (*garK, garR, garL, garP*) in the same transcriptional unit (*garP-garL-garR-garK-rnpB*) does not produce a fitness defect because essential gene expression can be rescued by the intra-operonic promoter (bottom panel). Targeting the intra-operonic promoter (rnpBp) or essential gene (*rnpB*) itself does produce a fitness defect. Each dot represents an sgRNA (centered at midpoint of chromosomal target) targeting either an essential (red-orange) or non-essential (gray) gene. **d** Fraction of non-essential genes upstream of an essential gene within the same transcriptional unit (TU) that also show a fitness defect when knocked down, likely indicating a CRISPRi-mediated polar operon effect.

We also measured the tightness of inducible control for the CRISPRi library by growing it with no inducer (i.e. no aTc or arabinose added to turn on expression of *dCas9* and sgRNA) for the same period of time as a regular fitness experiment (24 population doublings). Strains with essential gene-targeting sgRNAs exhibited a negligible growth defect in this uninduced condition (with gene fitness scores near 0), and the fitness defect of essential gene strains was significantly different between this uninduced case and an induced case (p < 0.001, Mann-Whitney U-test; Cohen’s d effect size = 3.7) (Fig. 2b). This suggested that library strains with sgRNAs targeting essential genomic features can be maintained when the library is propagated in an uninduced state.

We also checked if fitness was biased by factors such as position of targeting relative to chromosomal origin, GC content of the sgRNA, or chromosomal strand of the targeted gene and found no significant correlation (Supplementary Fig. 3). In agreement with prior reports of CRISPRi in bacteria^2,4-6^, we found CRISPRi-mediated polar operon effects where knockdown of an upstream nonessential gene in an essential gene containing operon produced a growth defect similar to the essential gene itself, indicating that CRISPRi can knockdown entire operons (Fig. 2c). Out of 160 operons containing at least one essential gene targeted in our library, we focused on 47 operons where the essential gene was not the first gene in the operon to assess the prevalence of polar operon effects. We found operon effects to be highly prevalent, with every non-essential gene (based on PEC database) upstream of the essential gene in 38 out of the 47 operons exhibiting a growth defect when targeted with dCas9 (Fig. 2d).

### CRISPRi Screening of Essential Genes Under Various Environmental Conditions

To evaluate whether CRISPRi could assess feature fitness in a condition-specific manner, we compared feature enrichment in the library by varying two physiologically relevant parameters – nutrient availability and oxygen availability. In the case of nutrient availability, we profiled the CRISPRi library in M9 media, M9 media supplemented with casamino acids (M9Ca), and LB media under aerobic growth conditions. In the case of oxygen availability, we profiled the CRISPRi library in LB media under aerobic and anaerobic growth conditions.

We first compared enrichment between varied nutrient availability conditions (LB, M9Ca, M9). As previously discussed, we saw a strong depletion of sgRNAs targeting known essential genes (based on knockout studies) in LB and M9 media. We next analyzed non-essential genes that should exhibit condition-dependent phenotypes between these conditions by comparing the enrichment of known amino acid metabolism genes for expected auxotrophic phenotypes. We found a strong depletion of guides targeting genes involved in amino acid biosynthesis in the amino acid-deficient medium (M9) but not the supplemented medium (M9Ca), indicating that CRISPRi can enrich for conditionally essential genes (Supplementary Fig. 4).

Finally, we looked beyond phenotypes for protein-coding genes and analyzed sRNA feature enrichment. Out of the 130 sRNAs with designed guides in the library, we had fitness data for 114 in each condition (some sRNAs did not have data due to low read depth in one or more conditions). Of these 114 sRNAs, we found novel phenotypes for the *hok/sok* Type I toxin-antitoxin (TA) system, which has been implicated in bacterial persistence through the stringent response^31,32^. Specifically, under stress or amino acid starvation, (p)pGpp and Obg induce (via an unknown mechanism) expression of the *hokB* toxin gene, which leads to membrane depolarization and persistence^33^. In our CRISPRi screens, a knockdown of the *sokB* antitoxin sRNA gene resulted in a successively stronger growth defect in LB, M9Ca, and M9 media (Supplementary Fig. 5), likely due to its inability to inactivate the *hokB* toxin gene product under conditions where it is expressed. The related *hokC-sokC* system exhibited a similar, yet even stronger, response to the knockdown of antitoxin *sokC*. Previous literature has suggested that *hokC* is likely inactive due to an insertion element located 22 bp downstream of the *hokC* reading frame^34^. However, the *sokC* antitoxin sRNA exhibits a strong deleterious phenotype when knocked down, implying that *hokC* may still be functional. We hypothesize that this phenotype was not seen earlier because the *hokC-sokC* system had only been investigated in nutrient-rich conditions (e.g. LB); however, here we are able to uncover this phenotype by combining the programmability of CRISPRi targeting to investigate this small 55 bp feature with the ability to assess feature fitness across conditions.

We next compared enrichment between the aerobically varied conditions, expecting to find condition-specific phenotypes for genes involved in aerobic or anaerobic growth processes. Many strains with guides targeting genes involved in aerobic respiration (eg pyruvate conversion genes, heme biosynthetic genes, ubiquinol biosynthetic genes, cytochrome *bd-I* terminal oxidase subunits, ATP synthase F_1_ synthase complex subunits) were depleted in the aerobic condition but dispensable under anaerobic growth (Fig. 3a). NADH:quinone oxidoreductase I (*nuoABCEFGHIJKLMN;* NDH-1) and NADH:quinone oxidoreductase II (*ndh*; NDH-2) showed a previously unreported phenotype (Supplementary Fig. 6). NDH-1 only exhibited a defect in aerobic minimal media conditions (M9Ca, M9) while NDH-2 only exhibited a defect in the aerobic rich media condition (LB), implying that NDH-1 may be the dominant oxidoreductase in nutrient-limited conditions and NDH-2 may be dominant in nutrient-rich conditions. We noted that seven genes (*hemB, hemC, hemD, hemH, ispB, nrdA, nrdB*) previously characterized as essential according to the Keio database of essential genes in *E. coli* K-12 BW25113^12^ and the PEC database of essential genes in *E. coli* K-12 MG1655 were dispensable for growth under anaerobic conditions (Fig. 3a). These genes are involved in heme biosynthesis (*hemB, hemC, hemD, hemH*) and ubiquinol biosynthesis (*ispB*), which play critical roles in the aerobic electron transport chain. The essential genes *nrdA* and *nrdB*, which are involved in aerobic nucleotide metabolism^35,36^, were also dispensable under anaerobic growth. We clonally verified the conditional essentiality of *nrdA* and *hemB* by showing that we could generate viable strains with deletions of these genes under anaerobic conditions and that these deletion strains were not viable under aerobic conditions (Fig. 3b-c, Supplementary Table 3). By demonstrating that these “essential” genes are only conditionally essential, we show that they are not part of the core, essential genome but instead part of the growth-supporting, conditionally-essential genome. We also noted that of the genes with conditional phenotypes in Fig. 3a, 20 were genes (genes with double asterisks in Fig. 3a) for which a gene disruption mutant was not generated during a high-throughput transposon insertion screen using Rb-TnSeq due to the attempted construction of the mutants under a condition where the underlying genes were essential. We clonally verified one of these genes, *ubiD*, by showing that we could generate a viable deletion strain under the condition determined as permissive via the CRISPRi screen (Fig. 3b-c, Supplementary Table 3). This analysis presents a proof of concept for the use of two intertwined capabilities of CRISPRi screening – the ability to induce CRISPRi to interrogate features traditionally regarded as essential and the ability to probe feature essentiality across conditions – to delineate between the core, essential and accessory, conditionally-essential genome.

**Fig. 3.**
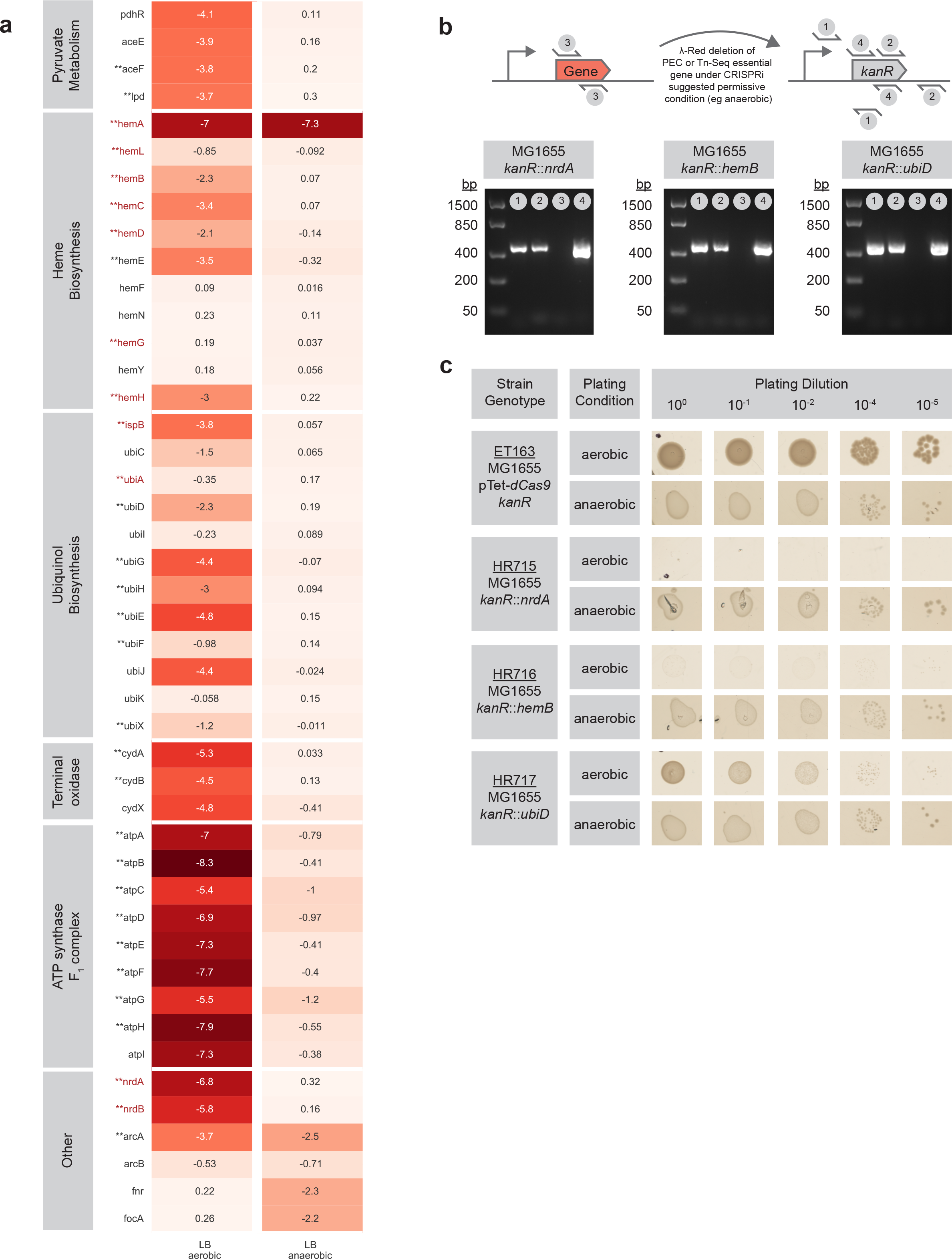
Conditional phenotypes from CRISPRi screening. **a** Comparison of CRISPRi phenotypes (gene fitness scores) between aerobic and anaerobic conditions in LB. Gene names in maroon represent genes classified as essential by the Keio collection (*E. coli* K-12 BW25113) and PEC database of essential genes in *E. coli* K-12 MG1655. Gene names with a preceding “**” superscript represent genes for which a mutant could not be generated using RbTnSeq during a high-throughput screen in *E. coli* K-12 BW25113. Gene fitness scores are averaged from a minimum of three replicates. **b** λ-Red recombineering mediated deletion of select aerobic essential genes from Keio collection/PEC database (*nrdA, hemB*) or sick genes from Rb-TnSeq (*ubiD*) under permissive condition (anaerobic) as discovered via the CRISPRi screen. Gel images with reactions validating in-frame deletion of each essential gene via PCRs showing successful integration of *kanR* resistance cassette and removal of essential gene at native gene locus. **c** Confirmation that anaerobically generated knockouts of selected genes are non-viable under aerobic condition (non-permissive condition). An MG1655 strain with *kanR* cassette integrated on the chromosome is provided as a WT-like reference (ET163).

### Time-series Measurements Elucidate Dynamic Knockdown Response to CRISPRi

We next leveraged the ability to induce CRISPRi perturbations on-demand to probe the dynamic response to knockdown for the library, focusing first on essential genes, and determine if time-series data could yield further measurement resolution into essential gene phenotypes. Specifically, we grew the induced library and sequenced samples at regular intervals over a period of 18 population doublings in LB rich media (Supplementary Fig. 7). We examined the fitness of strains harboring guides targeting essential genes across the timepoints and found that these strains exhibited successively stronger growth defects over progressive time points (Fig. 4a). We next clustered the essential gene time-series data (Supplementary Note 2) and found that essential genes could be classified into one of three groups (Early, Mid, Late) based on their temporal growth trajectory (see Figure 4b for examples and Figure 4c for groupings). For example, some essential genes showed a fitness defect soon after the first few population doublings while other genes did not show a defect until several population doublings had occurred. Of the 287 essential genes analyzed, 78 were in the Early group, 114 in the Mid group, and 95 in the Late group (Supplementary Table 4a).

**Fig. 4.**
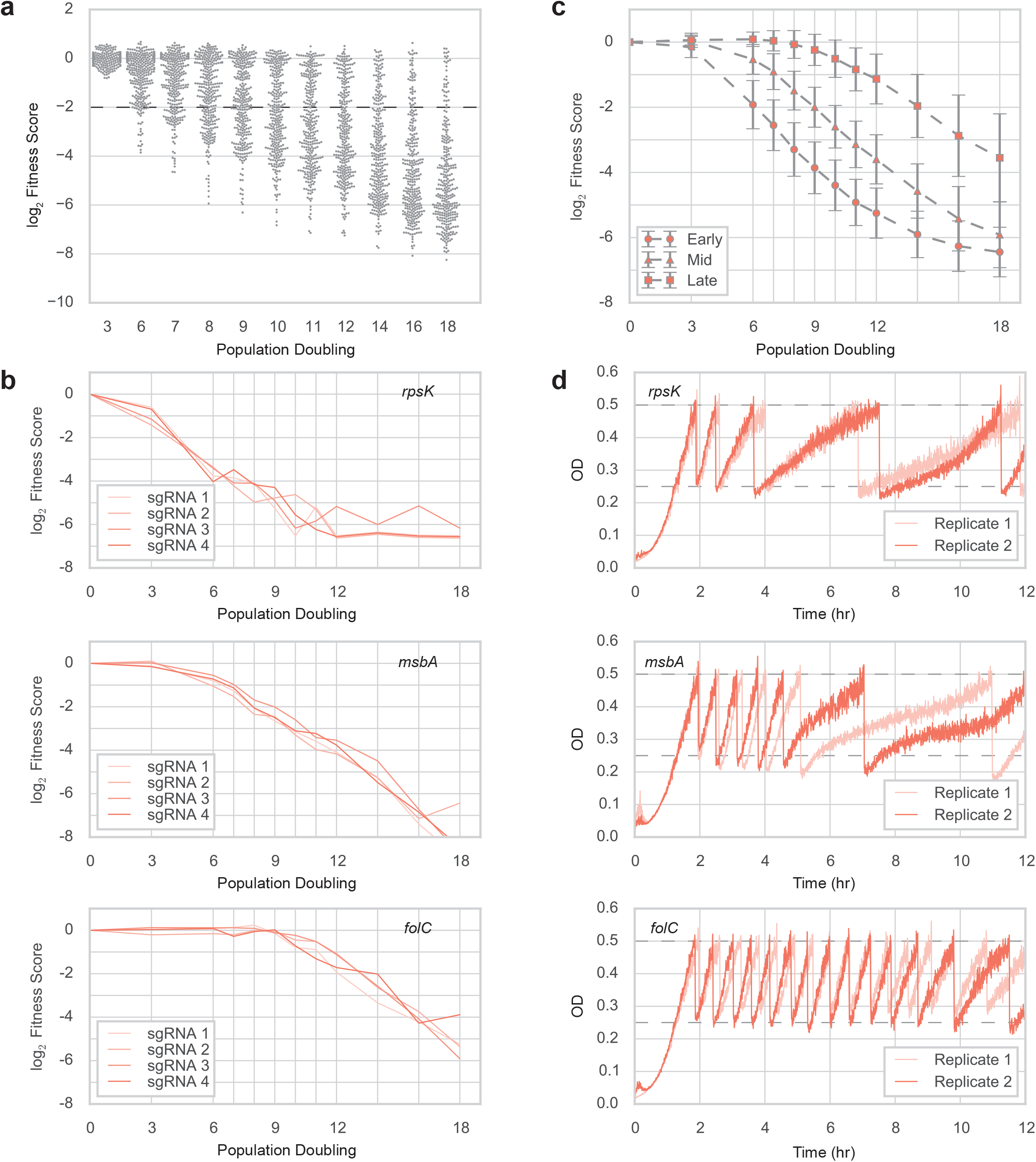
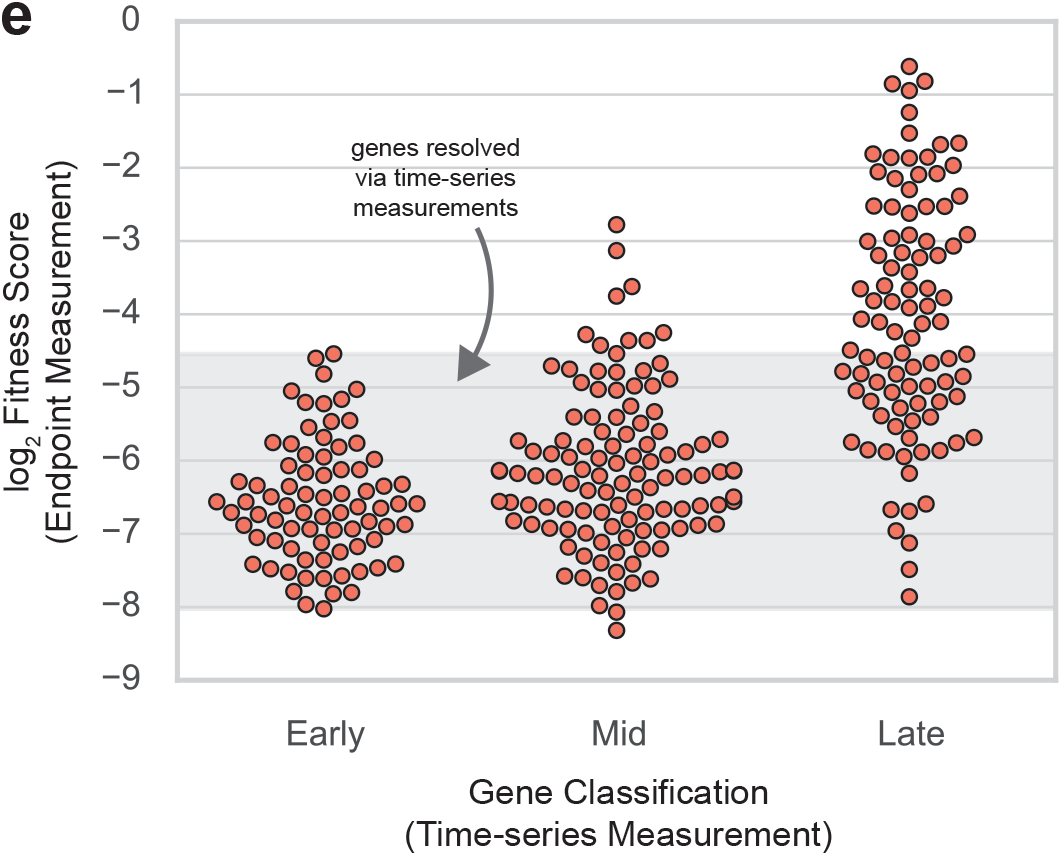
Temporal knockdown profiling of CRISPRi library. **a** Gene fitness scores for PEC essential genes (n=3O4) from pooled CRISPRi experiment calculated at progressive timepoints (e.g. population doubling 3, 6, 7...) relative to initial timepoint (T0). b Example temporal trajectories constructed from pooled CRISPRi experiment depicting one of three characteristic profiles observed for essential genes from K-means clustering. Each line represents an sgRNA for annotated gene (Early - *rpsK*, Mid - *msbA*, Late - *folC*). c Grouping of essential genes into classes (Early, Mid, Late) from K-means clustering and depiction of resulting composite growth curves. Each curve corresponds to an essential gene class with each solid marker (circle, triangle, square) denoting the mean fitness score of genes (averaged across two replicates) with that essential gene class at a given population doubling (n_Early_ = 78, n_mid_ = 114_nLate_ = 95; error bars represent ±1 standard deviation). d Growth curves of CRISPRi strains for candidate genes from each essential gene class as measured on eVOLVER, an automated turbidostat. For each selected essential gene, an sgRNA targeting that gene was selected from the CRISPRi library and cloned into a strain expressing dCas9. An uninduced culture of each strain was inoculated into the eVOLVER and grown until OD 0.50 in LB + antibiotics (carb/kan) media without inducers. Upon reaching this setpoint, each strain was diluted to OD 0.25 with LB + antibiotics (carb/kan) + inducers (aTc, arabinose) media and then allowed to grow between OD 0.25 and 0.50 with fresh inducer media being used for subsequent dilutions. Two replicates were grown for each CRISPRi gene strain. **e** Demonstration of additional resolution provided by time-series measurements (x-axis) in comparison to endpoint fitness measurements taken after 24 population doublings of the library (y-axis). Shaded region contains 219 essential genes for which stratification into gene classes provides additional phenotypic resolution.

We performed a gene ontology enrichment analysis to see if these classes were enriched for specific biological functions (Supplementary Table 4b, Supplementary Note 2). An analysis with TIGR Role ontologies^37^ revealed that essential genes in the Early group were significantly enriched for genes involved in ribosomal protein synthesis and modification (p < 0.001, p-value from Hypergeometric test followed by FDR correction) with 32 out of 41 essential genes with this TIGR Role present in the Early group. Resource allocation studies in *E. coli* have shown that in rapidly dividing cells ribosomes are most abundant and important for growth^38^ and haploinsufficiency studies in yeast have shown that ribosomal genes exhibit strong dose responses to gene expression perturbation in rich media^39^. This would support our finding of ribosomal protein synthesis and modification genes exhibiting a faster physiological response to expression knockdown (via growth defect) relative to other essential genes queried. An analysis of the Mid group revealed a strong enrichment in genes involved in tRNA aminoacylation (p < 0.001, Hypergeometric test with FDR correction) with 19 out of 22 essential genes with this TIGR Role present in the Mid group. The presence of tRNA aminoacylation genes in the Mid class also agrees with previous resource allocation studies, which report that the dosage effects observed under exponential growth are present, but less strong, for tRNA genes^40,41^. Finally, an analysis of the Late group revealed an enrichment of all eight essential genes involved in the 2-*C*-methyl-D-erythritol 4-phosphate/1-deoxy-D-xylulose 5-phosphate (MEP/DOXP) pathway (p < 0.05, Hypergeometric test with FDR correction). The MEP/DOXP pathway^42^ represents the mevalonate-independent pathway for producing the isoprenoid precursors isopentenyl pyrophosphate (IPP) and dimethylallyl pyrophosphate (DMAPP), and its presence in a later, albeit still essential class, in comparison to translation-related genes indicates that the abundance of certain pathway metabolites may not be as rate-limiting to growth in rich media as genes related to translation.

Because the transcriptional and translational products of genes expressed prior to essential gene knockdown are still present in cells upon CRISPRi induction, we hypothesized that differences in the initial abundance and decay rate of these products may affect the time it takes to observe a measurable fitness defect. We found that genes in the Early group had higher mRNA and protein abundance and were closer to the chromosomal origin than genes in other groups (Supplementary Fig. 8, Supplementary Table 4c); however, these trends were largely driven by the presence of protein synthesis related genes in the Early group and likely represent correlative effects as opposed to causative drivers of group classification.

We next analyzed all genes targeted in the library to see whether genes classified as non-essential also exhibited varied responses (Supplementary Note 2). We observed three categories after clustering, two of which contained genes exhibiting a growth defect via the knockdown of both essential and non-essential genes and a third category of genes that did not exhibit a growth defect (Supplementary Fig. 9, Supplementary Table 4d). Across the two categories of genes exhibiting a defect we saw an enrichment of a number of processes including translation, transcription, aerobic respiration, and fatty acid metabolism (Supplementary Table 4e).

The composite nature of the analyzed growth curves meant that the apparent decline in abundance of a given strain could be the result of the slower growth of that strain, the faster growth of another strain, or a combination of the two cases. To distinguish between these cases and validate the trends among essential gene classes, we chose a representative essential gene from each class (Early, Mid, Late), generated individual strains with dCas9 and sgRNAs to separately target these essential genes (Supplementary Table 5), and used the eVOLVER^43^, an automated cell culture system, to monitor the temporal knockdown response. We also generated a strain expressing dCas9 along with an sgRNA that did not target any genomic locus to serve as a reference control. We used the eVOLVER as a turbidostat by programming it to keep cells between two optical density (OD) ranges, which allowed us to track changes in doubling time in response to CRISPRi induction. The control sgRNA strain exhibited no change in doubling time after CRISPRi induction (Supplementary Fig. 10a). In comparing the essential gene-targeting validation strains, we found that *rpsK* (Early gene) was the first to show an increase in doubling time upon induction of CRISPRi, followed by *msbA* (Mid gene) and *folC* (Late gene), thus confirming our observations from the pooled screen (Fig. 4d). We also found that even within a gene class, different genes could have different profiles. For example, *msbA* showed a progressive increase in doubling time while *ftsZ* (another Mid gene) consistently showed a halt in cell growth after a set number of doublings (Supplementary Fig. 10b).

Having validated the trends among essential gene classes, we compared the time-series measurements from the screen to the endpoint measurements (i.e. fitness scores calculated from the initial and final time points in a screen) from an earlier screen. We found that the time-series measurements successfully enabled the further resolution of fitness for 219 essential genes via their stratification into gene classes (Figure 4e). Together, these results demonstrate (1) the increased measurement resolution provided by time-series measurements in resolving essential gene phenotypes and (2) that while CRISPRi knockdown of an essential gene eventually leads to a fitness defect, different genes can exhibit varied dynamic responses to perturbation, potentially indicating the functional importance of the genes and their biological roles as well as highlighting target considerations for CRISPRi applications where transient dynamics are important (e.g. CRISPRi-based genetic circuits).

### CRISPRi Screen Uncovers Design Considerations for Non-genic Targeting

*Promoter Interference:* The CRISPRi library contains 14,188 sgRNAs targeting 3,237 promoters and 4205 transcription start sites (TSSs) from RegulonDB (Supplementary Table 6a). To measure the efficacy of CRISPRi targeting for promoters on a genome-scale we assessed whether knockdowns of promoters regulating essential genes produced a growth defect (Fig. 5a). An analysis of 1,102 sgRNAs targeting 337 essential gene promoters across experiments in rich and minimal media (Supplementary Table 6b) revealed that (i) for 74% of essential gene promoters at least 1 sgRNA produced a mild knockdown phenotype (eg Fitness ≤ -1), and (b) for 51% of essential gene promoters, all sgRNAs produced a mild knockdown phenotype. Through this survey, we collected additional experimental phenotypes (i.e. a collection of fitness scores) for 141 known promoter annotations from RegulonDB, which primarily uses RNA-seq as the primary source of experimental characterization for promoters (Supplementary Table 7). We also found, to the best of our knowledge, the first phenotype-based experimental evidence for four computationally predicted promoters of essential genes (Supplementary Table 7), highlighting the utility of CRISPRi to improve the annotation strength of non-genic genomic features. We compared the fitness effect of targeting essential gene sequences to that of targeting promoter sequences of essential genes and found that targeting promoters to knockdown gene expression was less efficient than targeting the gene sequence itself (Fig. 5b). However, we did find cases where promoter-targeting produced a knockdown phenotype similar to gene-targeting knockdowns and where promoter-targeting yielded better knockdown performance than the gene knockdown (Supplementary Fig. 11), indicating the potential of promoter CRISPRi as an alternative to gene CRISPRi for control over gene expression. We also revisited the time-series data to analyze how promoter CRISPRi compared to gene CRISPRi following a perturbation. To avoid the confounding effects of multiple genes within the same transcriptional unit (TU) and multiple promoters driving the same TU, we focused on 27 monocistronic essential gene TUs regulated by a single promoter (Supplementary Note 2). We found a strong overlap between the trajectories of the two knockdown implementations (Supplementary Figure 12), which further indicated the potential of promoter CRISPRi in the presence of well-designed sgRNAs. To elucidate factors contributing to better promoter guide design, we analyzed cases where promoter CRISPRi failed. We noted that 91 % of essential promoters targeted by the 334 guides that did not produce a growth defect (Fitness > -1) either were part of a promoter array (i.e. two or more promoters in tandem regulating the same TU) or displayed a strong strand-dependency with respect to knockdown efficiency.

**Fig. 5.**
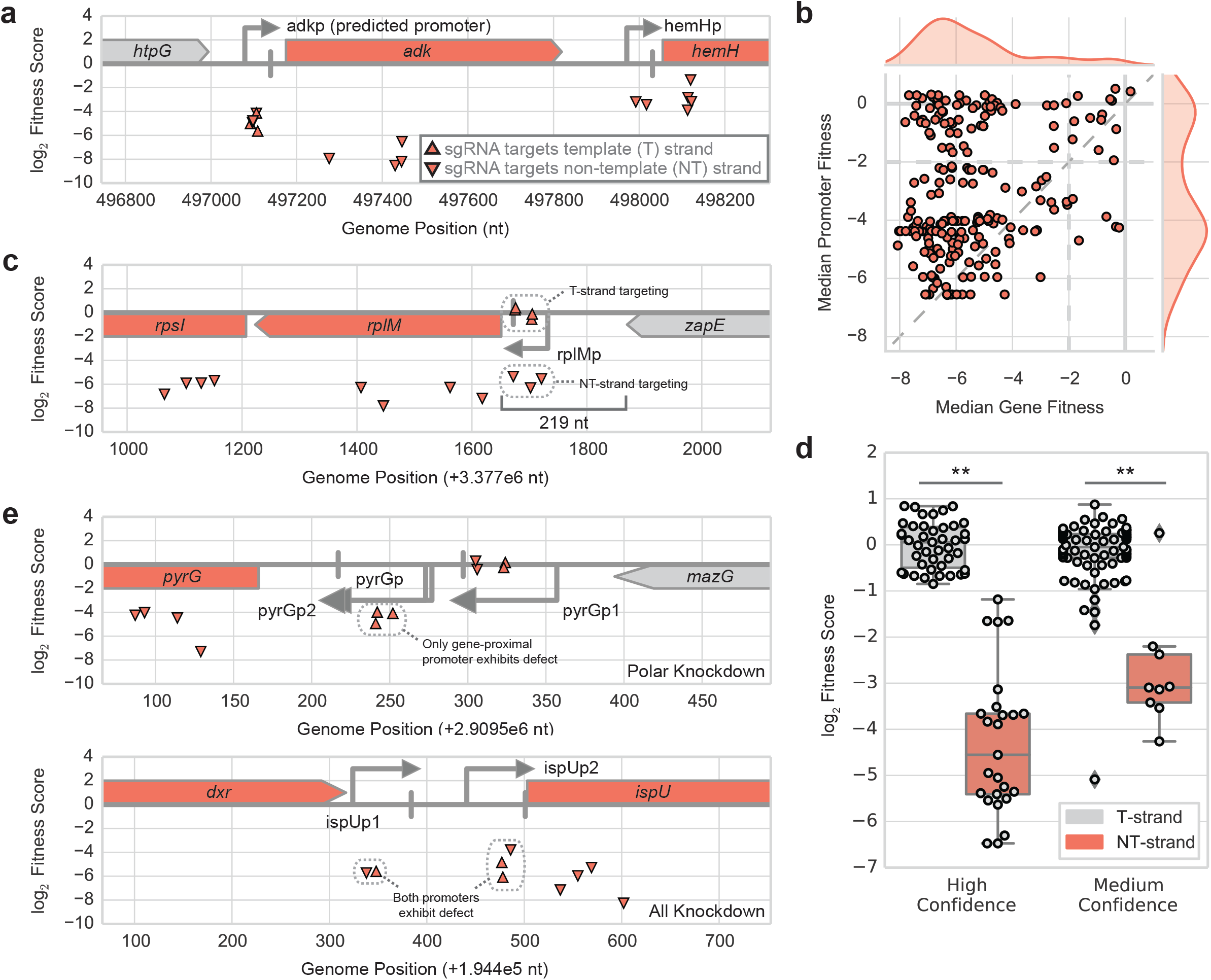
Non-genic phenotypes from CRISPRi library. **a** Demonstration of how CRISPRi fitness data for promoter knockdowns can add experimental confidence to predicted promoters (eg adkp) and known promoters (eg hemHp) by confirming that targeting the promoter produces a similar phenotype (i.e. fitness outcome) in comparison to targeting its regulated gene (eg adkp - *adk;* hemHp - *hemH*). **b** Comparison of efficacy of gene-targeting CRISPRi against promoter-targeting CRISPRi for gene expression knockdown. For all essential genes for which guides targeting both gene and promoter sequences were present in the library, the median of fitness scores for sgRNAs targeting the gene sequence (x-axis) is plotted against the median of fitness scores for sgRNAs targeting the promoter sequence (y-axis). Note that the thin diagonal dashed line represents y = x. **c** Depiction of strand-dependency of CRISPRi-mediated promoter knockdown for rplMp driving expression of the *rpIM-rpsI* operon. Only sgRNAs targeting the NT-strand of the promoter (relative to the gene) produce a fitness defect, while T-strand targeting sgRNAs do not. d Boxplots (with data points overlaid) showing strand dependent promoter CRISPRi for 12 high-confidence cases and 26 medium-confidence cases. Each case represents a TU and all of the promoters regulating it (Methods). **p < 0.001 (Mann-Whitney U-test); Cohen’s *d* = 4.3 (left), 3.2 (right). e Phenotypic profiles of tandem promoter arrays where only knockdown of an essential-gene proximal promoter yields a CRISPRi-mediated growth defect (top) or where a knockdown of any promoter regulating the essential gene can yield a growth defect (bottom).

We hypothesized that for cases where effective promoter knockdown was strongly dependent on the targeted strand, the sgRNAs could be targeting more effective positions within the promoter to interfere with transcriptional initiation or that the local genetic context was influencing knockdown efficacy. In the latter scenario, we hypothesized a model of “transcriptional coupling” where CRISPRi targeting of a promoter on the template strand failed to produce a fitness defect (while targeting the non-template strand could produce a defect) due to its inability to block RNAP readthrough from an upstream transcriptional event. We systematically identified 11 high-confidence cases where targeting the non-template strand produced a growth defect while targeting the template strand did not (Fig. 5c-d, see Supplementary Table 7 for cases and scoring metrics). One explanation for this result could be the transcriptional overlap of intra-operonic promoters in operons containing multiple TUs (e.g. one TU within a larger TU). Recent reports have also suggested that the transcription boundaries of operons are not as static as previously thought with one study using long-read sequencing (SMRT-Cappable-seq) to demonstrate that 34% of RegulonDB operons can be extended by at least one gene and that 40% of transcription termination sites have read-through that alters operon content^44^. Indeed, of the 11 high-confidence cases, five were TUs contained within larger operons and the remaining six TUs were a part of an extended RegulonDB operon in the SMRT-Cappable seq study (Supplementary Table 8). We also found an additional 26 cases of medium-confidence (Supplementary Table 7) that are candidates for this transcriptional coupling that we could not fully confirm either due to an insufficient number of guides available in both targeting orientations to test our strand hypothesis or due to cases where most, but not all, guides produced phenotypes matching the strand hypothesis (Fig. 5d, Supplementary Table 7). Of these 26 cases, 15 were TUs that were part of larger operons and seven were part of extended RegulonDB operons (Supplementary Table 8). Overall, our results suggest that targeting the non-template promoter strand can lead to a higher likelihood of successful CRISPRi knockdown for promoters in certain operonic contexts.

We also found that targeting CRISPRi in promoter arrays can yield distinct phenotypic profiles. Out of 59 tandem promoter arrays analyzed in the essential gene promoter data set, we found 40 tandem promoter arrays where we observed one of two distinct phenotypic profiles: (1) all promoters in the array produced a knockdown phenotype or (2) only the downstream promoter produced a fitness defect (Fig. 5e, Supplementary Table 7). In the case where all promoters produced a deleterious knockdown phenotype, we hypothesized that either the most upstream promoter was the primary driver of expression or that all promoters in the array were required for appropriate expression. In the case where only the downstream most promoter showed a deleterious knockdown phenotype, we hypothesized that either the downstream most promoter was the primary expression driver or that all promoters in the array are required for appropriate expression. The remaining 19 tandem promoter arrays analyzed either had an insufficient number of guides to draw any conclusions or were inconsistent with the aforementioned phenotypic profiles (Supplementary Table 7). Overall, our results showed that the promoter closest to the target gene is more likely to yield a knockdown phenotype and thus should be targeted when attempting to knockdown expression of a gene regulated by multiple promoters via promoter CRISPRi.

### TFBS Interference

Finally, we analyzed a set of 1810 sgRNAs in the library that were designed to target 1060 TFBSs on the chromosome (Supplementary Table 9). We first focused on a subset of 175 sgRNAs that targeted 102 TFBSs regulating an individual promoter controlling the expression of at least one rich media (based on PEC database) or minimal media (based on Joyce et al *J Bacteriol* 2006) essential gene. We found that most TFBS knockdowns that yielded a deleterious knockdown phenotype were present within the RNAP footprint for promoter binding, which we conservatively defined as between -60 to +20 nt relative to the transcription start site (TSS) associated with the promoter (Supplementary Fig. 13). Due to this overlap, we were unable to specifically associate such phenotypic outcomes to the TFBS alone as they could also be (and likely were) a result of promoter knockdown. Ultimately, we found that it was challenging to parse the phenotypic contribution of TFBSs due to their presence in promoters or binding site arrays with multiple diverse transcription factors.

We next looked at all TFBSs that exhibited a growth defect when targeted across all conditions in which the library was assayed. The activating NarL TFBS regulating the *cydDC* promoter, cydDp, exhibited a mild condition-dependent phenotype between aerobic and anaerobic conditions in LB (Supplementary Fig. 14). sgRNAs targeting *cydD*, which plays a role in respiration, and cydDp exhibited a growth defect in an aerobic fitness assay in LB medium but displayed no such defect under anaerobic conditions where no terminal electron acceptor was added and thus no respiration was active. Similarly, an sgRNA targeting the NarL TFBS, which has a positive effect on gene expression for *cydDC* and is situated -126 nt from the cydDp TSS, exhibited a mild growth defect as well (Fitness ~ -1.5) in the aerobic condition and a negligible growth defect (Fitness ~ 0) in the anaerobic condition.

## DISCUSSION

In this work we (1) explored key refinements to screen design and profiling using a genome-wide CRISPRi library in *E. coli* and (2) used the CRISPRi platform to phenotypically interrogate the *E. coli* genome. During the preparation of this manuscript, two other studies reported the use of genome-wide CRISPRi libraries to identify essential genes and genes involved in phage-host interactions in *E. coli*^4,45^. Our work here presents a complementary and extended demonstration of the power of CRISPRi-based approaches to interrogate microbial genomes with the discovery of novel phenotypes for essential genes using a more compact library, application of time-series measurements to track and elucidate phenotypic changes arising after CRISPRi induction, presentation of refined rules for CRISPRi targeting of promoters, and investigation of CRISPRi targeting of TFBSs.

We leveraged the inducible nature of CRISPRi to propagate strains with sgRNAs targeting essential genomic features and query them in a number of biochemical contexts, a task unfeasible using conventional gene disruption or knockout approaches. This enabled us to generate 100s of essential gene strains not covered by conventional knockout or Tn-Seq approaches in *E. coli*. Furthermore, we showed that a number of genes classified as essential genes according to classical aerobically generated *E. coli* knockout collections or unable to be assayed using Tn-Seq approaches were actually dispensable under anaerobic conditions, representing a more comprehensive annotation of these genes. We validated the dispensability of three of these genes by showing that we could generate strains with deletions of these genes under the condition they were predicted to be dispensable from the CRISPRi screen.

We also utilized the inducible nature of CRISPRi to track the effect of knockdown on essential genes post-induction of the CRISPRi machinery. Using time series measurements, we found that different essential gene strains displayed growth defects at distinctly different times, and our results enabled us to classify essential genes into specific categories based on how quickly a given gene’s knockdown yielded a measurable fitness defect. The genes in the most essential category had a remarkable overlap with genes discovered to be most essential in other resource allocation studies of *E. coli* in the same condition and also matched gene dosage studies in yeast. Overall, our results enabled us to further resolve the strong fitness defects associated with essential gene expression perturbation, and the use of time-series measurements in future high-throughput genetic screens should yield insight into the temporal importance of essential processes in conditions of interest.

The programmable nature of CRISPRi targeting also allowed us to interrogate promoters and TFBSs. Specifically, we were able to compare gene-targeted CRISPRi (inhibit transcription elongation) to promoter-targeted CRISPRi (inhibit transcription initiation), finding that gene-targeting CRISPRi largely outperformed promoter-targeting CRISPRi. We also attributed phenotypic evidence to 141 known RegulonDB-annotated promoters and associated, to our knowledge, the first experimental evidence to four predicted promoters from RegulonDB. Finally, we explored phenotypic profiles associated with tandem promoter arrays and promoters that displayed strand-dependent knockdown success to conclude that targeting the NT-strand of the promoter closest to the target gene can yield more successful CRISPRi knockdowns in comparison to other promoter-mediated orientations for certain genomic contexts.

While we demonstrated a high utility for microbial genome interrogation via CRISPRi-based screens in this work, CRISPRi still has a number of limitations. First, targeting in operons yields polar effects, thus limiting the analysis of essentiality to transcriptional units and assigning specific phenotypic confidence to only the last gene in the transcriptional unit. As such, CRISPRi should serve as a complementary method to transposon insertion and recombineering-based approaches, which are less prone to polar operon effects. Second, the compact organization of bacterial genomes yields architectures with overlapping or tightly spaced TFBS and promoter features. This makes it especially challenging to precisely attribute phenotypes to a specific TFBS (due to its proximity or overlap with other TFBSs and promoters). Precise genome editing methods such as MAGE and CREATE are likely more suitable for such cases. Regardless, the programmability of CRISPRi targeting can be used to uncover intergenic regions of phenotypic importance through tiled screens, which can be combined with TFBS and promoter predictions along with high-throughput measurements (eg protein-DNA interactions, RNA-seq) to add annotation confidence for newly-sequenced microbes. Overall, the CRISPRi library developed here presents a resource of curated and phenotype-linked sgRNAs for use in *E. coli*, and the workflow developed here for interrogating genic and non-genic chromosomal features provides the basis for high-throughput CRISPRi studies in other bacteria.

## METHODS

Chemicals, reagents, and media: LB Lennox Medium (EZMix™ powder microbial growth medium, Sigma Aldrich) was used to culture strains for experiments in rich media. M9 Minimal Medium (1X M9 salts, 2 mM MgSO_4_, 0.1 mM CaCl_2_, 0.4% glycerol) was used to culture strains for experiments in minimal media. Anhydrotetracycline (aTc; CAS 13803-65-1, Sigma-Aldrich) was used at 200 ng/mL to induce dCas9 expression. Arabinose was used at 0.1% to induce sgRNAs. Antibiotic concentrations used were 100μg/mL for carbenicillin and 30μg/mL for kanamycin. Glucose was used at 0.2% in media for outgrowth of the library from a freezer aliquot. Casamino acids (0.2%) were also used in M9 Minimal Medium for select assays.

### CRISPRi library design

See Supplementary Note 1 for details regarding sgRNA library design along with Supplementary Table 1 for sgRNA feature annotations, sequence-level details, and a summary of category codes.

### CRISPRi library construction

To clone the sgRNA library, sgRNAs were amplified from the OLS oligo pool using primers 282 (5’ CACATCCAGGTCTCTCCAT 3’) and 284 (5’ cacatccaggtctctCGGACTAGCCTTATTTTAACTTG 3’) using Phusion II HS and the following protocol: 98°C for 10 sec and 15 cycles of 98°C for 10 sec, 60°C for 30 sec, and 72°C for 20 sec followed by a final extension of 72°C for 5 min. The PCR reaction was purified using a Zymo DNA Clean & Concentrator kit and eluted in water. The purified library was cloned into the library receiver plasmid, pT154 (https://benchling.com/s/seq-YGEVpcmWzQjGfRrP8oDc), via a golden gate reaction using BsaI and T7 DNA ligase. The golden gate reaction product was purified using a Zymo DNA Clean & Concentrator kit, following the kit parameters for a plasmid cleanup. A derivative of *Escherichia coli* K-12 MG1655 (ET163: MG1655 FRT-*kanR*-FRT *tetR-pTet-dCas9;* https://benchling.com/s/seq-Gxu6IV96FF6y8jycpTrU) was used as the recipient strain for the sgRNA library. The purified library was electroporated into a competent cell preparation of ET163 and maintained under carbenicillin (plasmid marker) and kanamycin (strain marker) selection. Aliquots of the resulting library were stored at -80°C.

### CRISPRi fitness experiments

An aliquot of the library was taken from storage at -80°C and thawed at room temperature. The aliquot was used to inoculate a 5 mL culture of LB Lennox media (LB) with carbenicillin, kanamycin, and glucose (multiple aliquots were used to inoculate distinct cultures for experiments with biological replicates). The culture was grown at 37°C until it reached OD600 0.5. A 4 mL aliquot was taken as an initial time point for the library (t0 sample); this sample was centrifuged (Eppendorf 5810R) at 4000 RPM (3202xg) and stored at -80°C. The remaining 1 mL of culture was centrifuged (Eppendorf 5417R) at 8000xg and washed twice with 1 mL of LB media. 156 uL of this washed sample was added to 10 mL of LB media (~1:64 dilution) with arabinose (0.1%), aTc (200 ng/mL), carbenicillin (100μg/mL), and kanamycin (30μg/mL). Technical replicates were generated by dividing this initial culture into 5 mL cultures. Cultures were grown at 37°C until they reached OD600 ~0.5, indicating 6 population doublings of the library. The library was again diluted 1:64 into 5 mL of LB media with arabinose, aTc, carbenicillin, and kanamycin and grown at 37°C until the culture reached OD600 ~0.5. This process was repeated until the library had undergone a total of 24 population doublings under induction. After 24 population doublings, the sample was centrifuged (Eppendorf 5810R) at 4000 RPM (3202xg) and stored at -80°C.

For experiments in minimal media, the original freezer aliquot of the library was inoculated in M9 media with glycerol (0.4%), glucose (0.2%), carbenicillin, and kanamycin. For induction of the CRISPRi system, the library was cultured in M9 media with glycerol, arabinose, aTc, carbenicillin, and kanamycin. Casamino acids (0.2%) were added depending on the assay condition.

For time-series experiments, samples were collected every doubling after the t0 sample was taken for the first 12 doublings, after which samples were collected every two doublings until the library had undergone a total of 18 doublings. During the experiment, the library was maintained between OD600 ~0.25 and ~0.50.

### CRISPRi sequencing library preparation

Frozen, centrifuged samples from fitness experiments were taken from storage at -80°C and thawed at room temperature. The CRISPRi sgRNA library was isolated using a QIAprep® Spin Miniprep Kit. 10-20 ng of DNA from each sample was used for a PCR reaction to generate NGS-ready sequencing samples in a 50 uL reaction using Phusion polymerase and two primers to add one of two sets of indexed Illumina adaptors. The first set contained a constant reverse primer and a variable forward primer with sample-specific 8 nucleotide barcodes that were sequenced “in-line” during an Illumina sequencing read. The second primer set contained a constant forward primer and a variable reverse primer with sample-specific indices that could be sequenced during an indexing read (Supplementary Table 1d). Both primer sets yielded comparable sequencing results; however, we eventually shifted to using the second primer set as the data could be readily demultiplexed using Illumina software.

Each reaction was performed using the following protocol: 98°C for 30 sec and 21 cycles of 98°C for 10 sec, 67°C for 15 sec, and 72°C for 10 sec followed by a final extension of 72°C for 5 min. 5 uL of each PCR sample was pooled and purified using a Zymo DNA Clean & Concentrator kit. The purified sample was quantified using the Qubit dsDNA HS assay kit and product size was confirmed using a Bioanalyzer 2100 automated electrophoresis system (DNA 1000 Kit). Final samples were run on either an Illumina Miseq or HiSeq instrument (2000/2500; Vincent J. Coates Genomics Sequencing Laboratory, UC Berkeley). All relevant sequencing data have been deposited in the National Institutes of Health (NIH) Sequencing Read Archive (SRA) at https://www.ncbi.nlm.nih.gov/bioproiect/PRJNA559958 under Accession code PRJNA559958.

### CRISPRi sequencing data analysis

Sequencing runs were demultiplexed using standard Illumina software for samples using the second primer set or a custom python script (demultiplex_fastq.py) for samples using the first primer set. Demultiplexed reads were processed using the following set of custom python scripts: trim_sgRNA_reads.py to trim and filter reads according to quality thresholds; bwa_samtools.py to map the trimmed sgRNA reads to a BWA index of the sgRNA library; parse_bam.py to convert mapped reads to a table of counts that represent the abundance of each sgRNA in the sample. Custom scripts for analysis are available at https://github.com/hsrishi/HT-CRISPRi.

### CRISPRi fitness score calculation

A small constant (i.e. pseudocount of 1) was added to the raw read counts to avoid errors in calculating fold-change in subsequent fitness calculations due to division by 0. These adjusted read counts for each sample were normalized by the median abundance for that sample, thus generating relative abundance (RA) values for each sgRNA library member and enabling comparisons between different samples. The fitness score was calculated as the log2 ratio of the RA of a guide strain in a test condition relative to its RA in a control condition. In this framework, the test condition was a sample of the library after being subjected to grown over the course of an experiment, and the control condition was the t0 sample. The fitness scores from each sample were normalized such that the median fitness score for the sample was 0. In practice, library members with t0 raw read counts < 10 were filtered out to limit variability due to low read depth. Significance values for each sgRNA fitness score were calculated via the edgeR package using raw read counts as the input^46,47^.

We also created a gene fitness score, which we calculated as the median of fitness values for all sgRNAs targeting a given gene. This provided a more stringent metric for quantifying strong fitness scores. For example, for a given gene with four sgRNAs, at least two guides would have to yield a strong fitness score in order for the median to be lower than -2. Fitness scores for all relevant experimental samples are listed in Supplementary Table 10.

## AUTHOR CONTRIBUTIONS

H.S.R. led the experimental work and computational analyses. H.S.R, E.T., and A.P.A designed experiments. E.T. cloned the CRISPRi library and performed initial experiments. H.L. and X.W. designed the sgRNA library. A.P.A. supervised the research. All authors contributed to manuscript preparation. L.S.Q. and A.P.A. conceived of the research.

## ACKNOWLEDGEMENTS

The authors would like to thank (1) Agilent Technologies for providing the sgRNA library, (2) members of the Arkin lab, especially Vivek Mutalik and Morgan Price, for insightful discussions over the course of the work and during manuscript preparation, and (3) Guillaume Cambray and David Chen for initial help with Illumina NGS read processing. This work used the Vincent J. Coates Genomics Sequencing Laboratory at UC Berkeley. The authors would also like to acknowledge funding sources: Department of Energy Genome Science program - Office of Biological and Environmental Research [DE-SC008812, Funding Opportunity Announcement DE-FOA-0000640]; National Science Foundation (NSF) Graduate Research Fellowship (to H. S.R.); National Institutes of Health (NIH) Genomics and Computational Biology Training Program [5T32HG000047-18] (to H.S.R.). Funding for open access charge: DOE BER [DE-SC008812].

## Supplementary Information

### Table of Contents

**Supplementary Figures**

- Supplementary Figure 1: CRISPRi library replicability
- Supplementary Figure 2: CRISPRi library minimal media experiment
- Supplementary Figure 3: Investigation of bias in CRISPRi library
- Supplementary Figure 4: CRISPRi library amino acid auxotrophy experiment
- Supplementary Figure 5: Conditional phenotypes for *hok-sok* toxin-antitoxin system
- Supplementary Figure 6: Conditional phenotypes for NADH:quinone oxidoreductases
- Supplementary Figure 7: Workflow for CRISPRi time-series experiment
- Supplementary Figure 8: Analysis of gene product features on time-series gene classification
- Supplementary Figure 9: Time-series classification of all genes in CRISPRi library
- Supplementary Figure 10: eVOLVER profiling of ctrl and *ftsZ* CRISPRi strains
- Supplementary Figure 11: Examples of promoter-targeting guides more effective than genetargeting guides
- Supplementary Figure 12: Comparison of promoter- and gene-targeting CRISPRi time series
- Supplementary Figure 13: CRISPRi knockdown of TFBSs regulating single essential gene promoters
- Supplementary Figure 14: Feature cofitness of *cydD* gene, promoter, and TFBS-targeting sgRNAs

**Supplementary Tables**

- Supplementary Table 1: CRISPRi library design details (separate attachment)
- Supplementary Table 2: List of genes with median gene fitness score > -2 (separate attachment)
- Supplementary Table 3: List of essential gene knockout validation strains (this file)
- Supplementary Table 4: Gene classification and ontological enrichment from time-series analyses (separate attachment)
- Supplementary Table 5: List of strains used for eVOLVER CRISPRi experiment (this file)
- Supplementary Table 6: Annotations for sgRNAs targeting promoters (separate attachment)
- Supplementary Table 7: Results from analysis of essential gene promoters (separate attachment)
- Supplementary Table 8: Comparison of transcription readthrough results with SMRT-Cappable Seq study (separate attachment)
- Supplementary Table 9: Annotations for sgRNAs targeting TFBSs (separate attachment)
- Supplementary Table 10: Fitness scores for relevant experimental samples (separate attachment)

**Supplementary Notes**

- Supplementary Note 1: sgRNA library design
- Supplementary Note 2: Analysis of time-series data

**References (SI only)**

---

**Supplementary Fig. 1.**
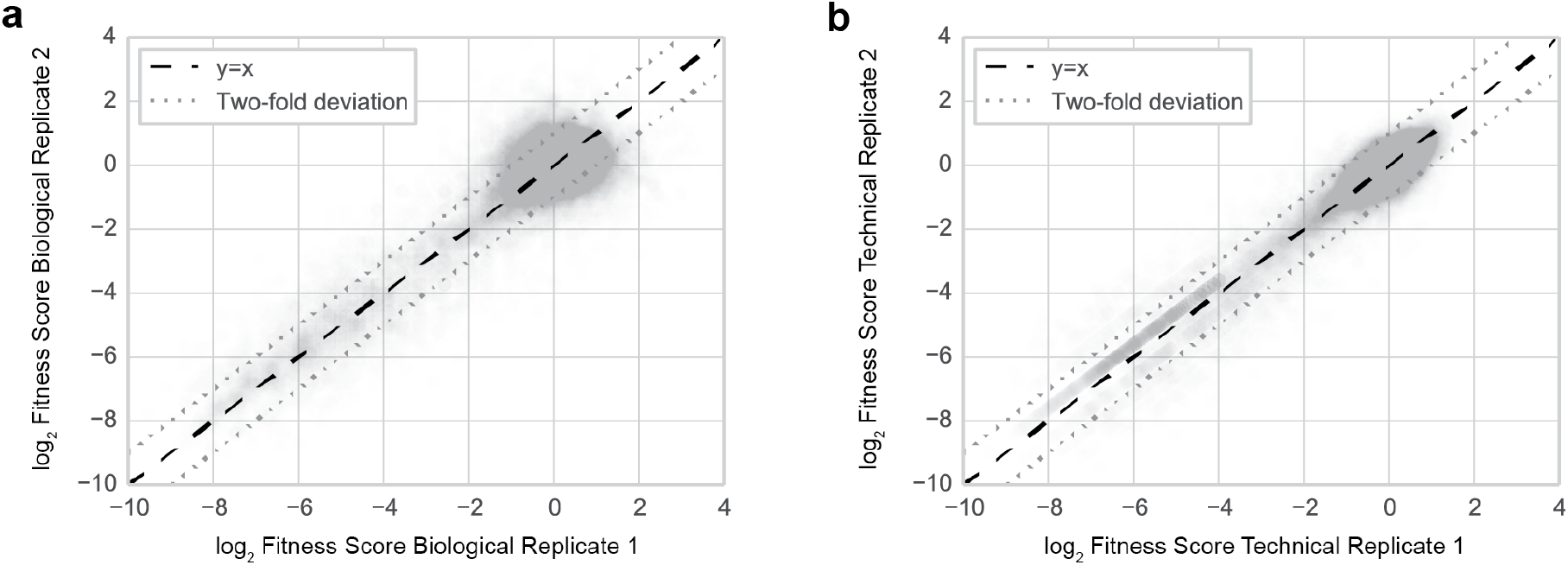
CRISPRi library replicability. **a** Two biological replicates of a CRISPRi experiment where the library was grown in LB rich media. Each dot represents an sgRNA. A biological replicate represents a distinct library aliquot, b Two technical replicates of a CRISPRi experiment where the library was grown in LB rich media. Each dot represents an sgRNA. A technical replicate represents an aliquot of the library that was split prior to the start of the experiment.

**Supplementary Fig. 2.**
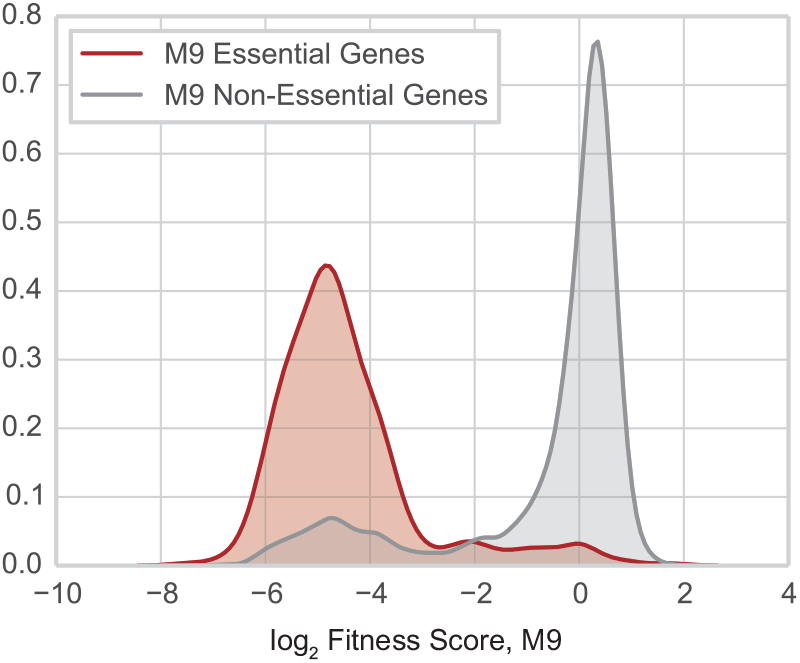
CRISPRi library minimal media experiment. Depletion of minimal media (M9) essential gene targeting sgRNAs compared to non-essential gene targeting sgRNAs over the course of a pooled fitness experiment with the HT-CRISPRi library in M9 minimal media (with CRISPRi system induced) under aerobic growth conditions for 24 population doublings.

**Supplementary Fig. 3.**
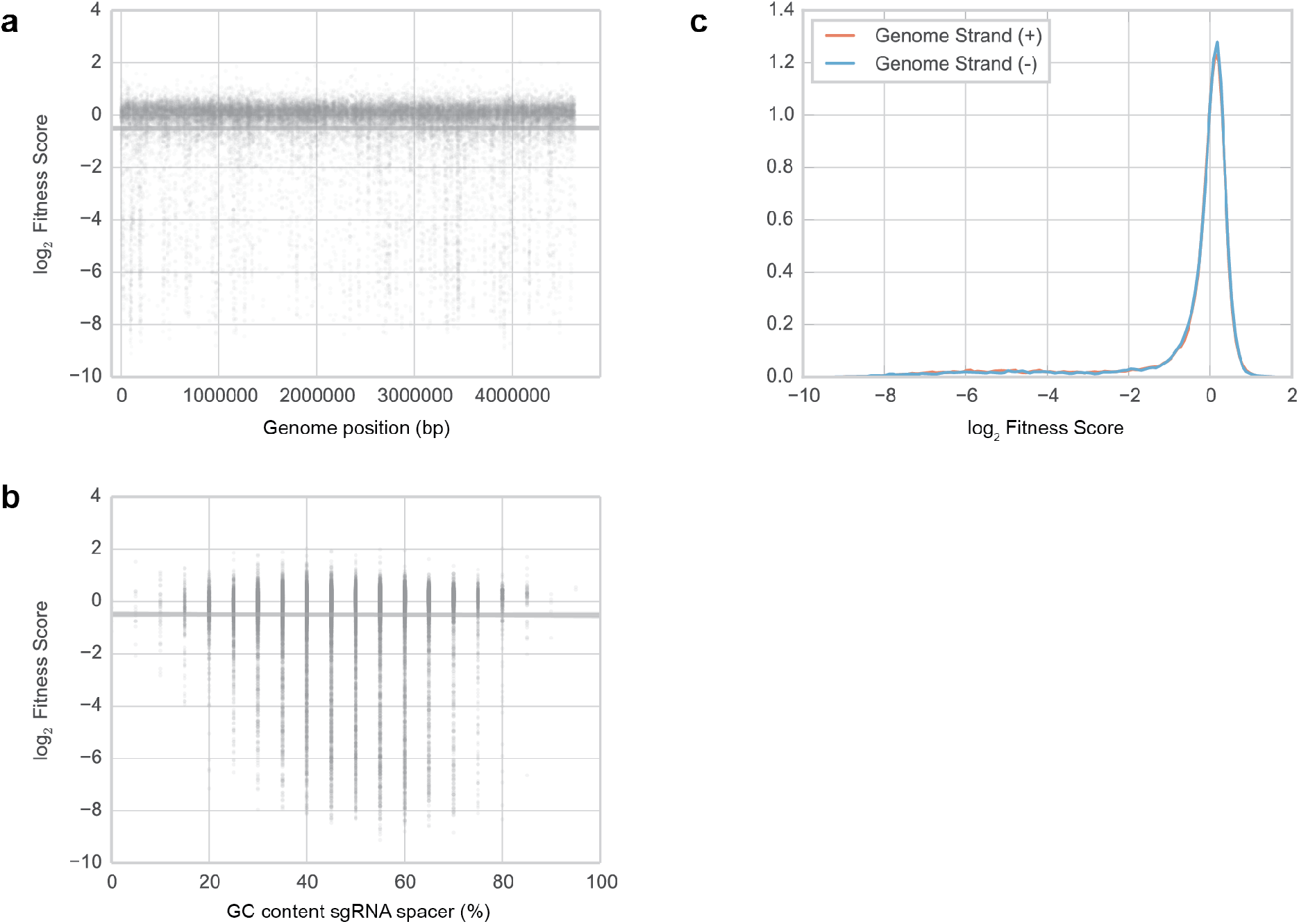
Investigation of bias in CRISPRi library. **a** Genome position of library sgRNAs plotted against fitness of respective sgRNAs from a pooled experiment in LB media under aerobic conditions. Gray line represents linear relationship between fitness and genome position with 95% confidence interval, b GC content of sgRNA variable region for library sgRNAs plotted against fitness of respective sgRNAs from a pooled experiment in LB media under aerobic conditions. Gray line represents linear relationship between fitness and GC content of sgRNA spacer with 95% confidence interval, c Distribution of fitness scores for sgRNAs targeting features on the + or - strand of the genome.

**Supplementary Fig. 4.**
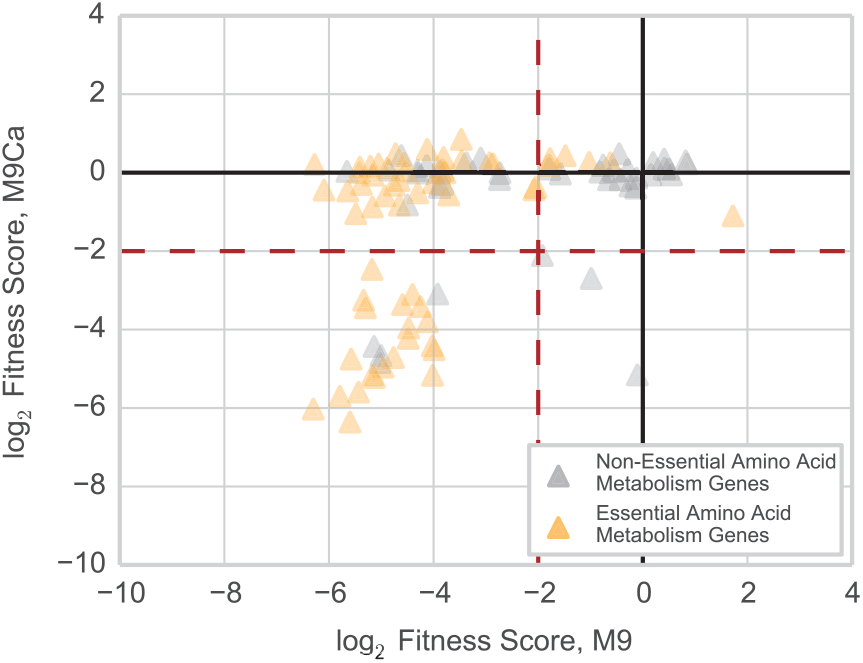
CRISPRi library amino acid auxotrophy experiment. Depletion of amino acid biosynthetic gene targeting sgRNAs over the course of a pooled fitness experiment in either M9 minimal media (x-axis - M9) or M9 minimal media supplemented with casamino acids (y-axis - M9Ca) under aerobic growth conditions for 24 population doublings. Essential amino acid metabolism genes (yellow triangles) refer to genes classified as essential in Joyce et al *J Bacteriol* 2006 via screening of the Keio essential gene deletion collection on glycerol minimal medium.

**Supplementary Fig. 5.**
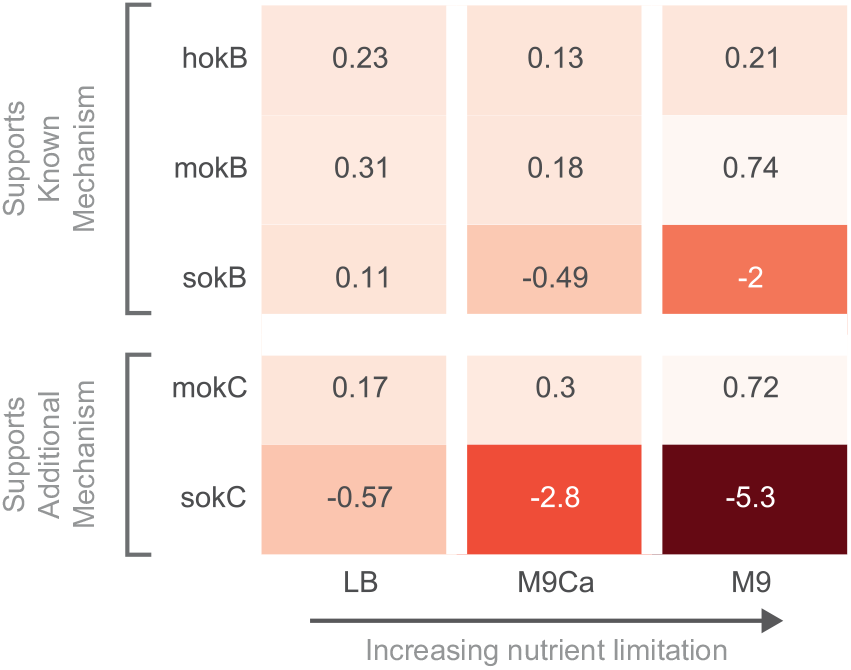
Conditional phenotypes for *hok-sok* toxin-antitoxin system. Gene fitness scores for genes in the *hok-sok* toxin-antitoxin systems (B & C) showing increasing defect as a result of *sokB* and *sokC* knockdown under conditions of increasing nutrient limitation with *sokC* depicting a stronger phenotypic response than *sokB*. Mechanism for hokB-sokB is reported in Verstraeten et al *Molecular Cell* 2015. Nutrient conditions: LB (rich media), M9Ca (M9 minimal media supplemented with casamino acids), M9 (M9 minimal media). Gene fitness scores are averaged from a minimum of three replicates. Data from pooled fitness experiment with library grown for 24 population doublings under induction in stated condition.

**Supplementary Fig. 6.**
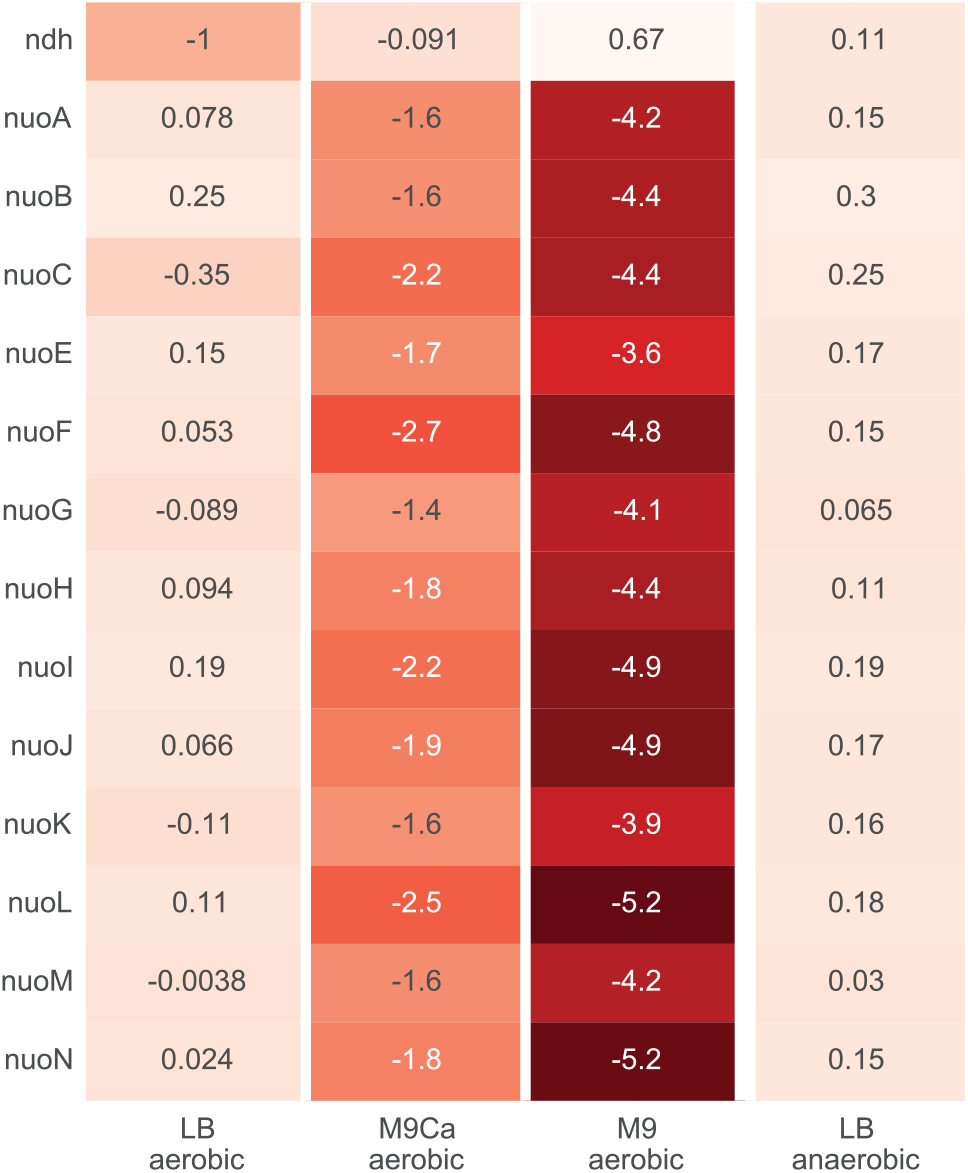
Conditional phenotypes for NADH:quinone oxidoreductases. Comparison of CRISPRi phenotypes (gene fitness scores) between aerobic conditions in LB, M9Ca, and M9 media against anaerobic condition in LB for NADH:quinone oxidoreductase I (NDH-1; *nuo* genes) and NADH:quinone oxidoreductase 2 (NDH-II; *ndh*). Gene fitness scores are averaged from a minimum of three replicates. Data from pooled fitness experiment with library grown for 24 population doublings under induction in stated condition.

**Supplementary Fig. 7.**
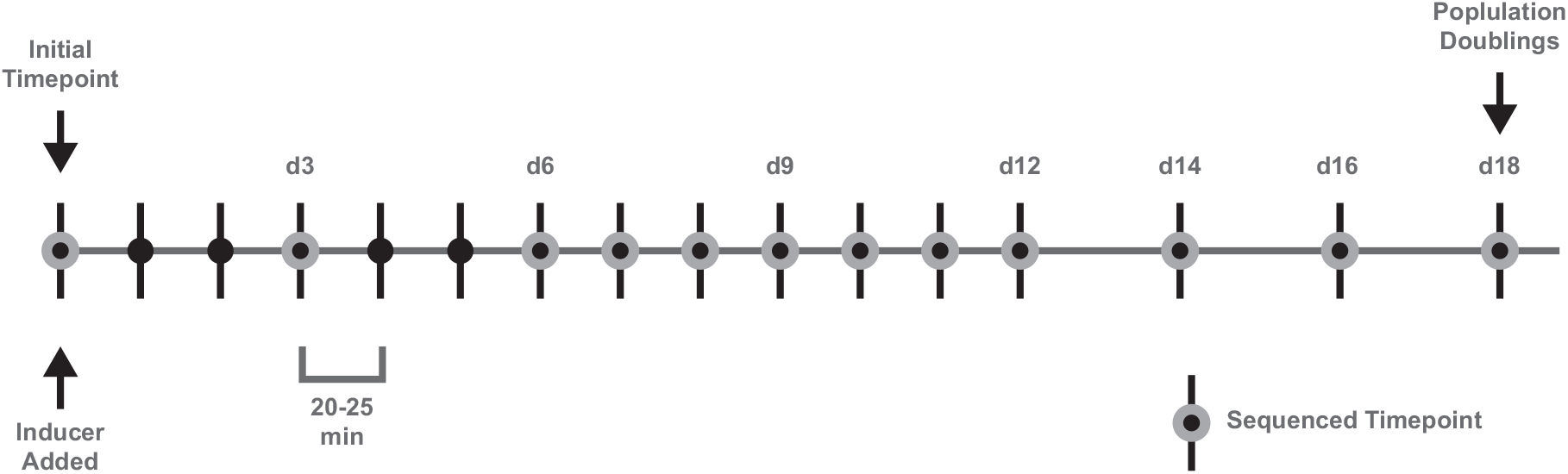
Workflow of CRISPRi time-series experiment. The library was induced and an initial timepoint was taken. Samples of the library were taken every population doubling for the first 12 doublings and then every other doubling until population doubling 18. Timepoints with gray circles were sequenced.

**Supplementary Fig. 8.**
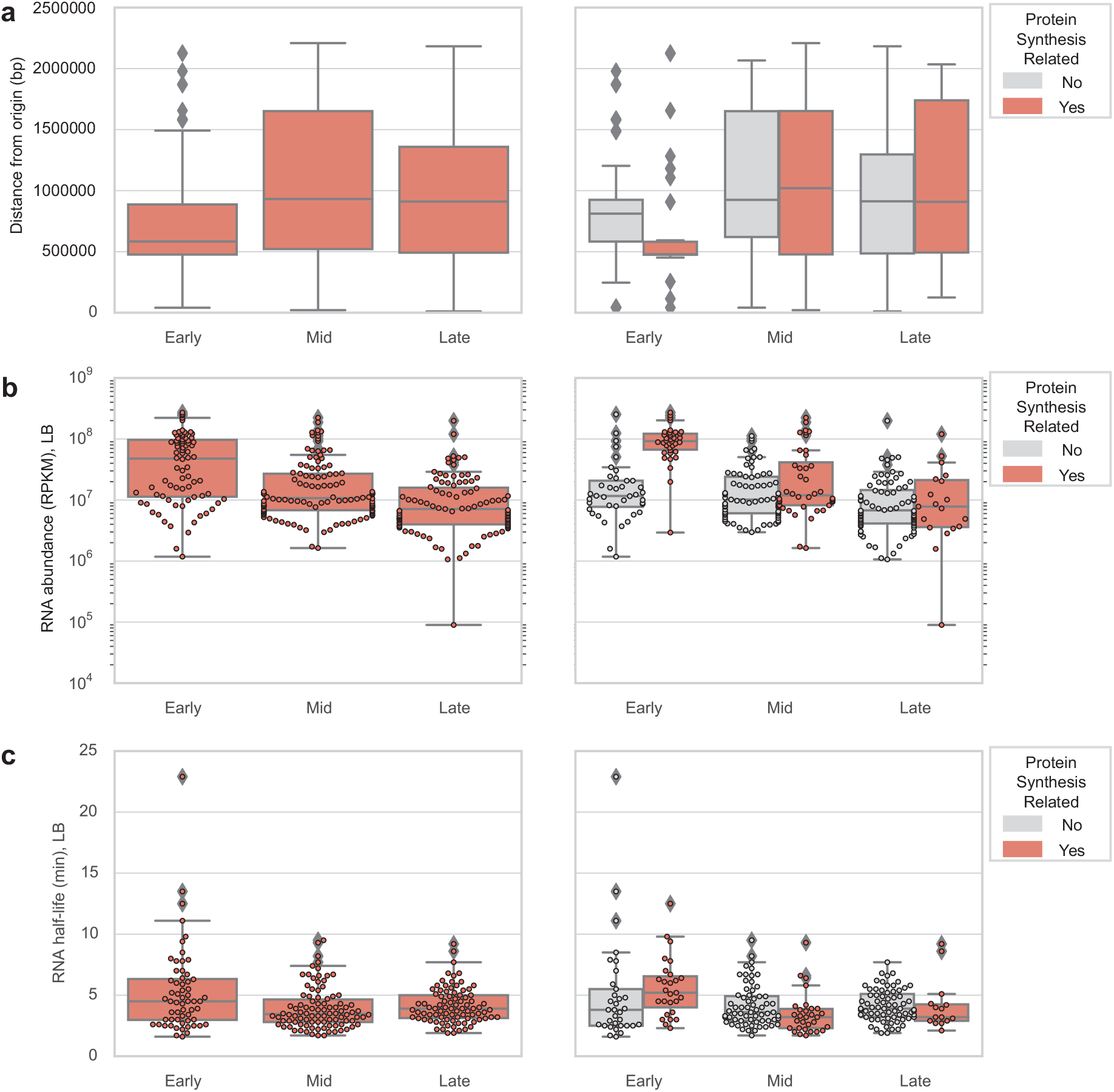
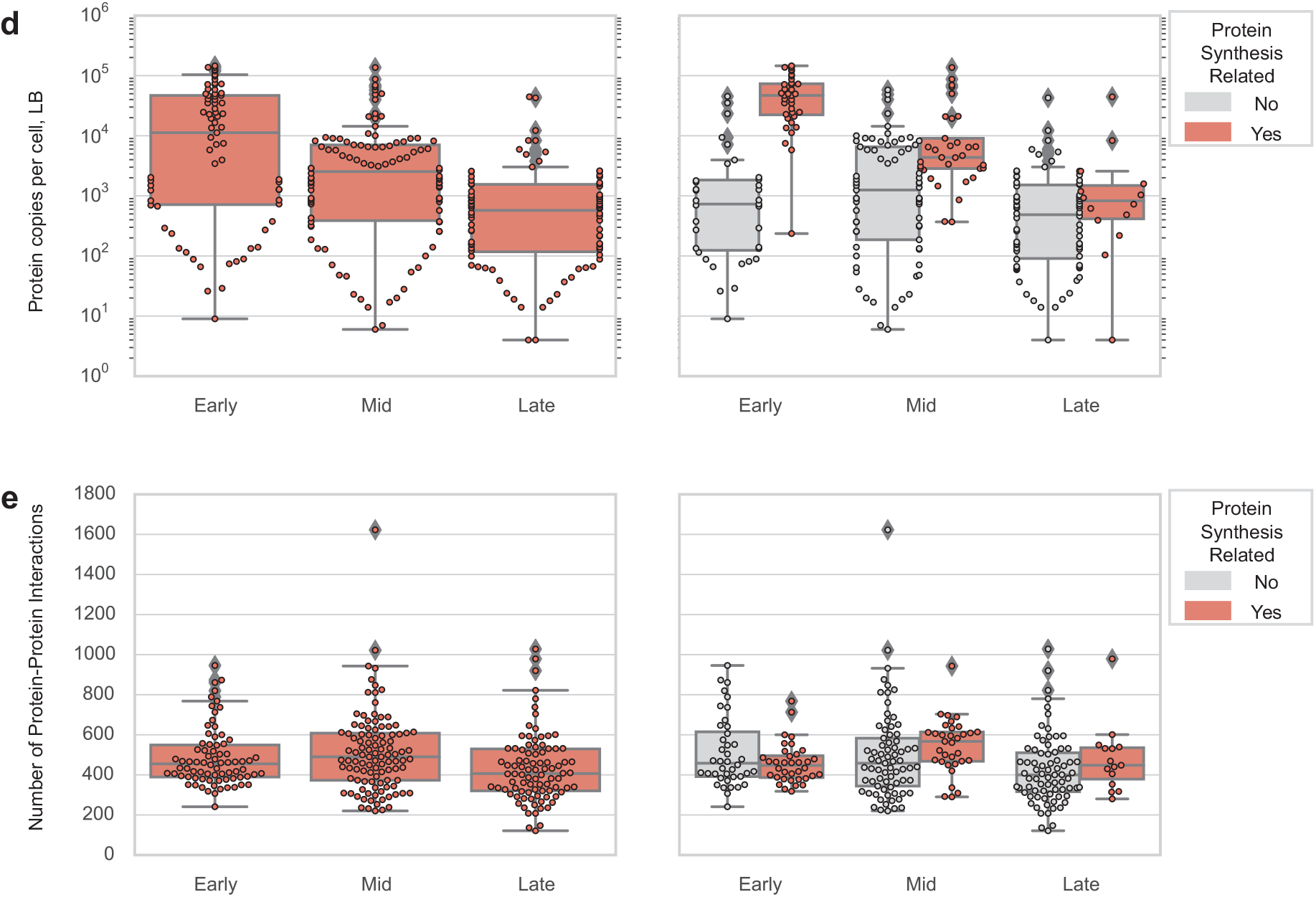
Analysis of gene product features on time-series gene classification. Comparison of effect of (**a**) distance of gene start position from origin of replication, (**b**) RNA abundance in LB media, (**c**) RNA half-life in LB media, (**d**) protein abundance, and (**e**) number of protein-protein interactions for genes in each essential gene class from time-series data. Essential gene class comparisons (left) are further decomposed into subgroups of genes related (or not) to protein synthesis (right; protein synthesis related defined as corresponding TIGR Roles with leading descriptor “Protein synthesis” - e.g. “Protein synthesis:tRNA aminoacylation”) to show if these genes are drivers of class-level trends. mRNA half-life data sourced from Venturelli et al *Nat. Comm*. 2017 GEO accession GSE94998. mRNA half-life data sourced from Bernstein et al *PNAS* 2002 Table 5. Protein abundance data sourced from Schmidt et al *Nat Biotechnol* 2016 Table S6. Protein-protein interaction data sourced from STRING database (string-db.org v10.5).

**Supplementary Fig. 9.**
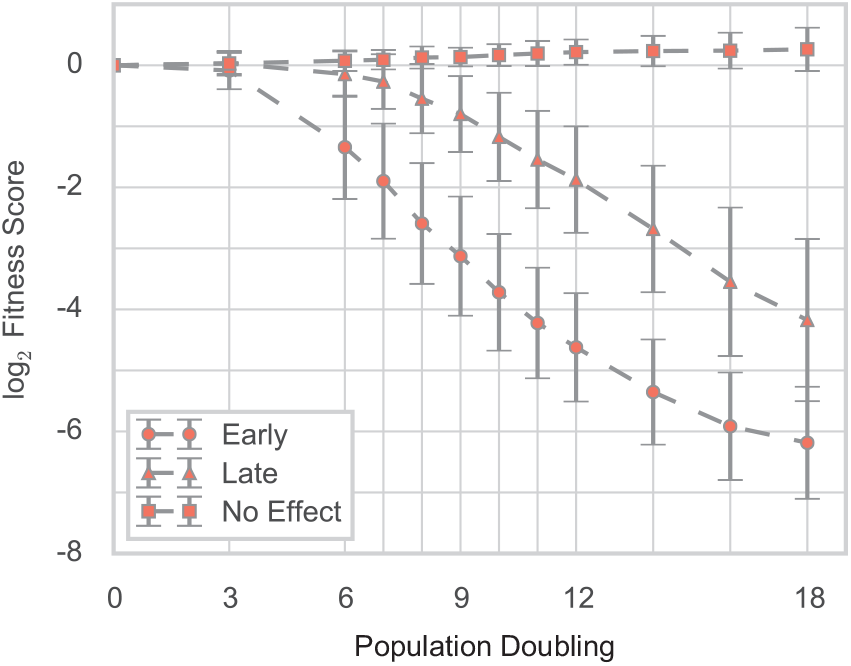
Time-series classification of all genes in CRISPRi library. Grouping of all genes targeted in CRISPRi library into classes (Early, Late, No Effect) from K-means clustering and depiction of resulting composite growth curves. Each curve represents a gene class with each solid marker (circle, triangle, square) denoting the mean fitness score of genes (averaged across two replicates) with that gene class at a given population doubling (n_Early_ = 188, n_Late_ = 218, n_No Effect_ = 4046; error bars represent ±1 standard deviation).

**Supplementary Fig. 10.**
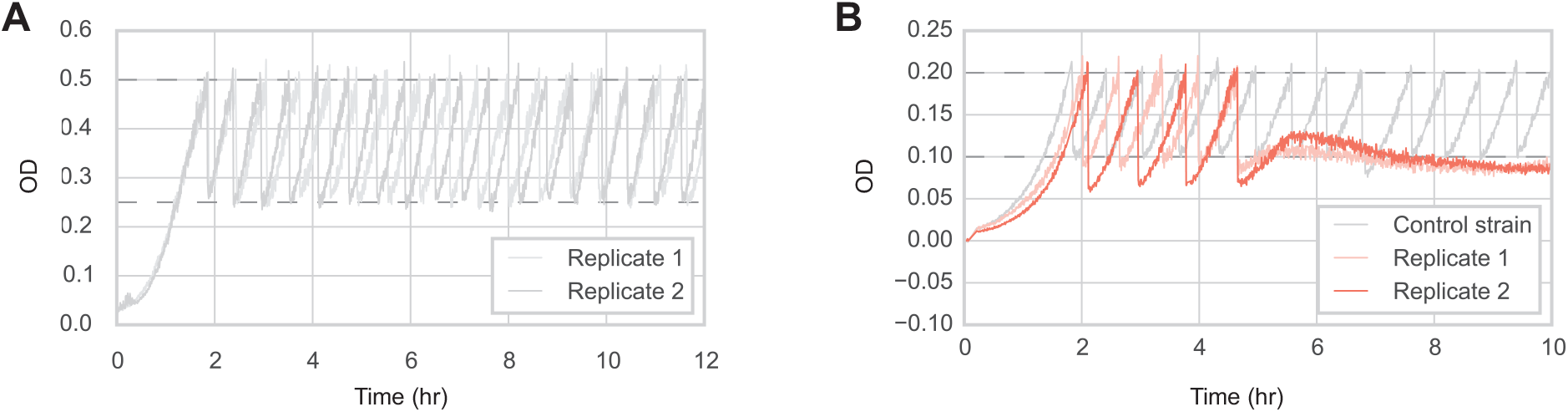
eVOLVER profiling of control and *ftsZ* CRISPRi strains. **a** eVOLVER growth curves of two replicates of a CRISPRi strain expressing dCas9 and a control sgRNA that does not target any locus on the chromosome. An uninduced culture of the strain was inoculated into the eVOLVER and grown until OD 0.50 in LB + antibiotics (carb/kan) media without inducers, after which each strain was diluted down to OD 0.25 with LB + antibiotics (carb/kan) + inducers (aTc, arabinose) media and then allowed to grow between OD 0.25 and 0.50. **b** eVOLVER growth curves of replicate *ftsZ*-targeting CRISPRi strains. An sgRNA targeting *ftsZ* was selected from the CRISPRi library and cloned into a strain expressing dCas9. An sgRNA designed to not target any locus in the *E. coli* genome was also cloned into a strain expressing dCas9 and used as a reference control strain. An uninduced culture of each strain was separately inoculated into the eVOLVER and grown until OD 0.20 in LB + antibiotics (carb/kan) media without inducers, after which each strain was diluted down to OD 0.10 with LB + antibiotics (carb/kan) + inducers (aTc, arabinose) media and then allowed to grow between OD 0.10 and 0.20 for multiple generations until ~10 hours.

**Supplementary Fig. 11.**
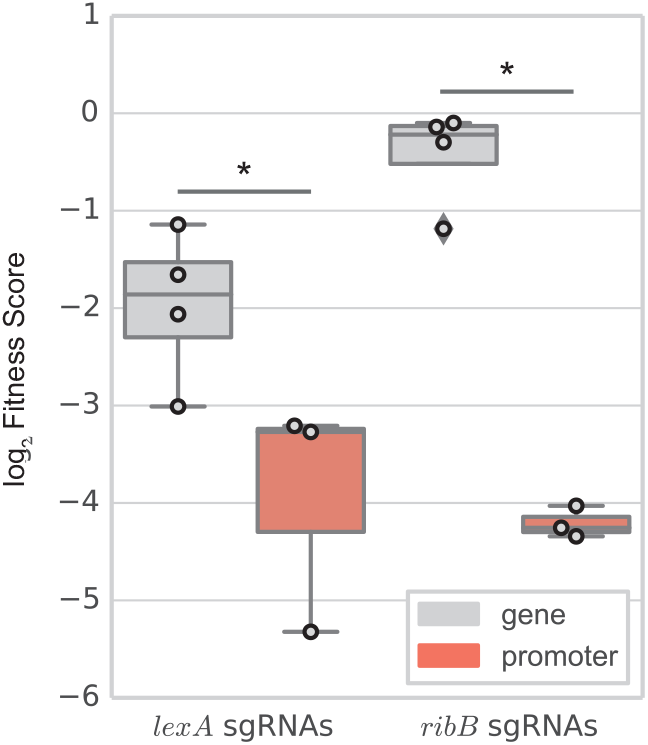
Examples of promoter-tageting guides more effective than gene-targeting guides. Example case where promoter-targeting sgRNAs provide better knockdown of a known essential gene than functional gene-targeting sgRNAs (left - *lexA*) and gene-targeting sgRNAs that were unable to produce a fitness defect (right - *ribB*). *p < 0.05 (Mann-Whitney U-test); Cohen’s *d* = 2.4 (left), 10.7 (right).

**Supplementary Fig. 12.**
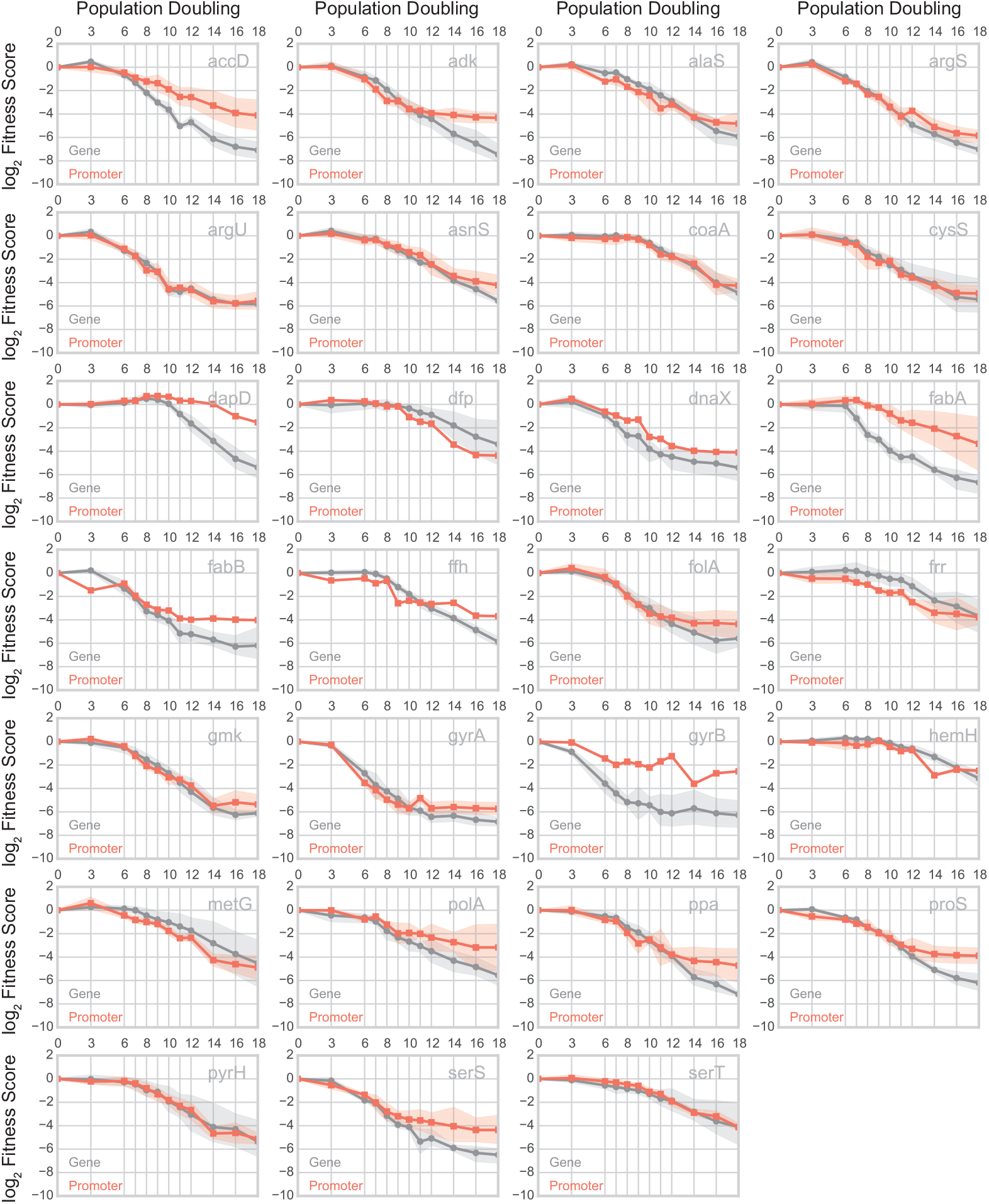
Comparison of promoter- and gene-targeting CRISPRi time series. Composite fitness curves of promoter- and gene-targeting sgRNAs with Fitness ≤ -1 for monocistronic essential gene transcriptional units regulated by a single promoter (see Supplementary Note 2 for details). Each curve represents the mean fitness of gene- (gray; circle marker) or promoter- (red-orange; square marker) targeting sgRNAs (averaged across two replicates) for each measured time point with corresponding shaded regions repressenting 95% confidence intervals.

**Supplementary Fig. 13.**
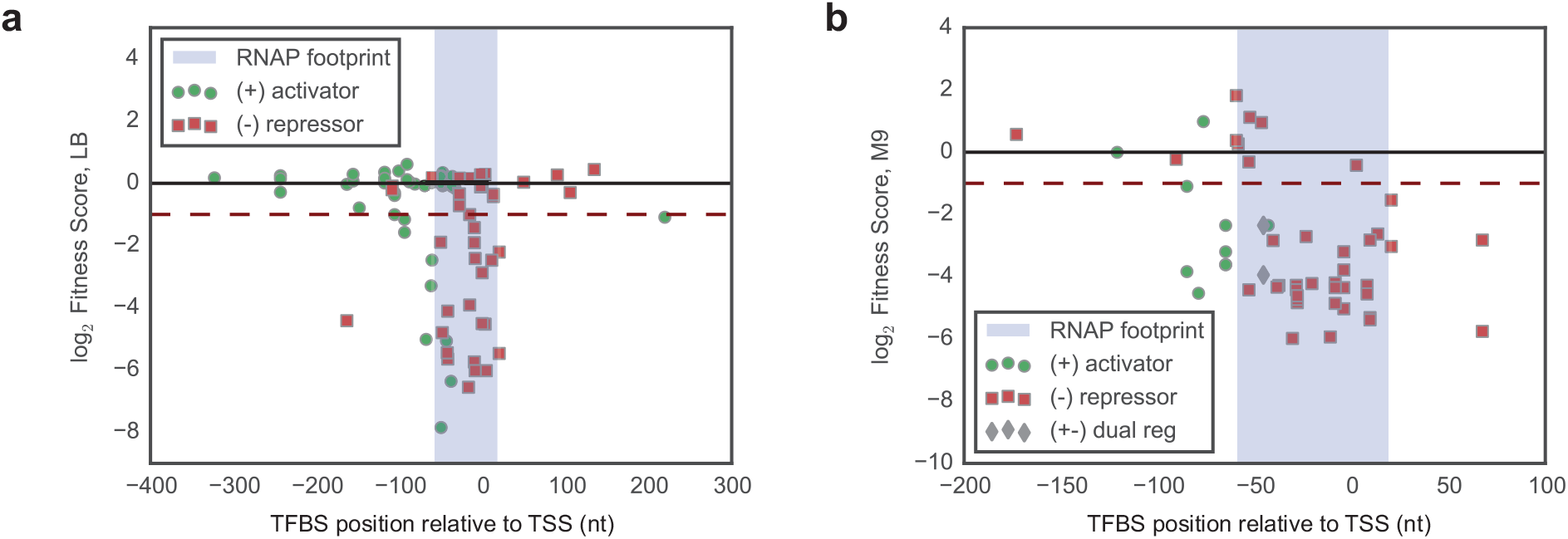
CRISPRi knockdown of TFBSs regulating single essential gene promoters. **a** Fitness scores for sgRNAs targeting TFBSs regulating single promoters of transcription units containing at least one LB essential gene (as determined by PEC database). The RNAP footprint is defined as the window between -60 to +20 nt relative to the transcription start site (TSS) of the regulated promoter. Each object in the scatter plot represents the fitness of an sgRNA (y-axis) targeting a TFBS at a given distance from the TSS of the promoter it regulates (x-axis). A given TFBS can have a positive effect on gene expression (green circles), negative effect on gene expression (red squares), or dual effect on gene expression (gray diamonds) as determined by RegulonDB annotations. **b** Fitness scores for sgRNAs targeting TFBSs regulating single promoters of transcription units containing at least one M9 essential gene (as determined by Joyce et al *J Bacteriol* 2006).

**Supplementary Fig. 14.**
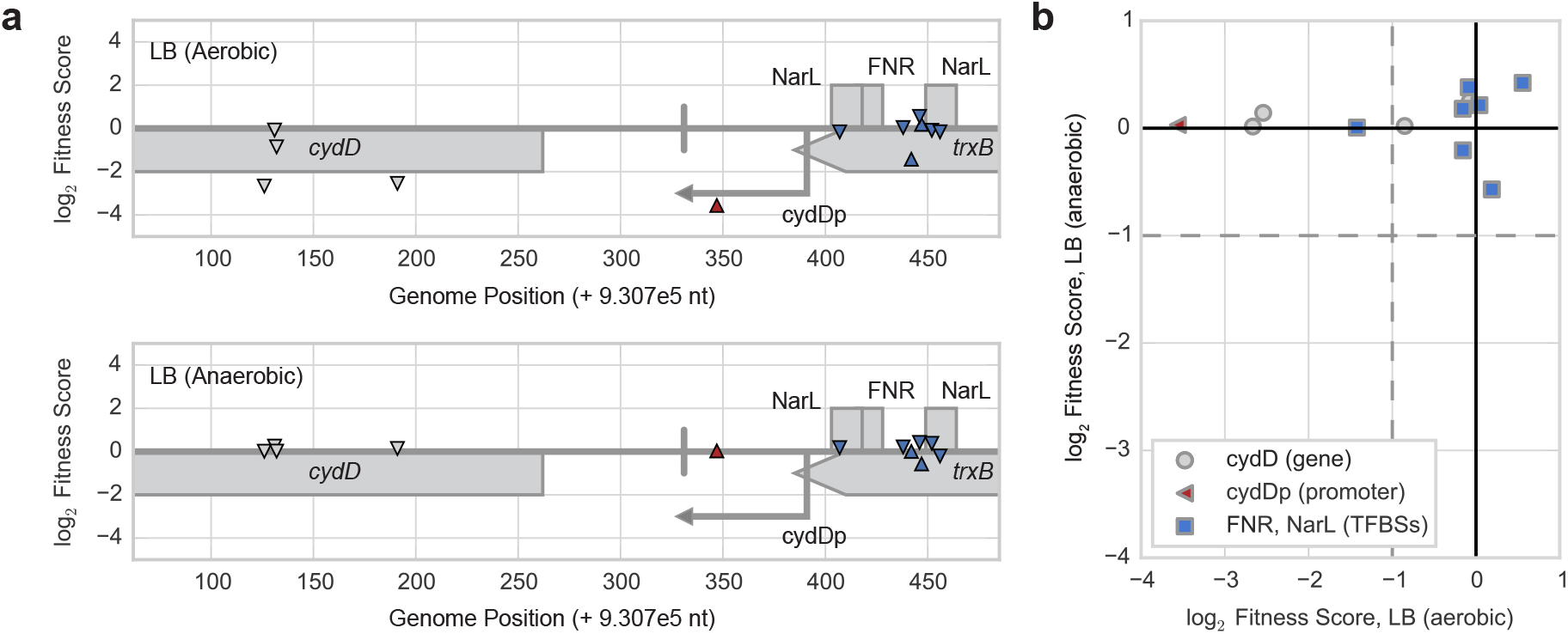
Feature cofitness of *cydD* gene, promoter, and TFBS-targeting sgRNAs. **a** Fitness data for *cydD* gene, its corresponding promoter (cydDp), and TFBSs (NarL - gene expression activator, FNR - gene expression activator) regulating its promoter from fitness assays in LB media between aerobic (top panel) and anaerobic (bottom panel) conditions. Each triangle represents an sgRNA (centered at midpoint of chromosomal target) targeting either the chromosomal strand corresponding to the non-template (downward facing triangle) or template (upward facing triangle) strand of the *cydD* gene. **b** Scatter plot comparing conditional phenotypes for sgRNAs targeting *cydD* (gray circles), *cydD* promoter (red triangle), and *cydD* TFBSs (blue squares) between aerobic and anaerobic conditions.

**Supplementary Table 3.**
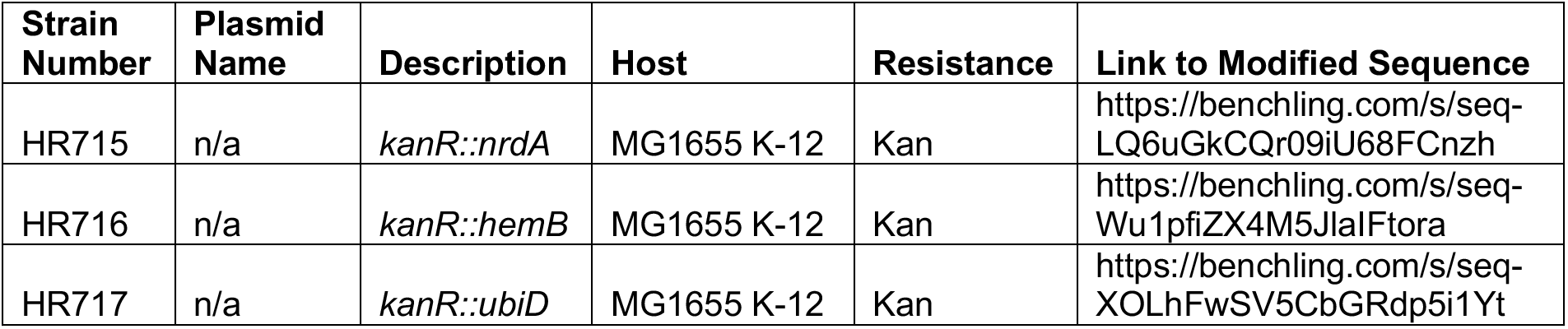
List of essential gene knockout validation strains.

**Supplementary Table 5.**
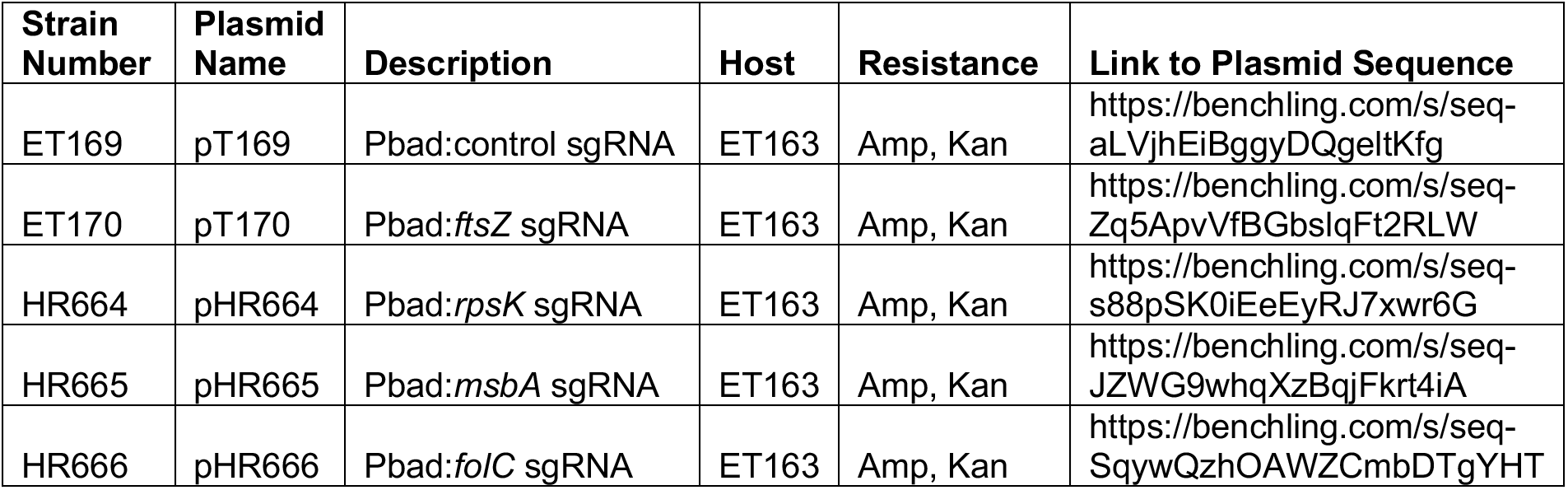
List of strains used for eVOLVER CRISPRi experiment.

## Supplementary Note 1. sgRNA library design

We designed the sgRNA library following rules described in Larson et al *Nature Protocols* 2013:

Selection of sgRNAs for oligo pool:

1. We first identified all 5’- XXXXX XXXXX XXXXX XXXXX NGG-3’ sequences by searching both the sense and anti-sense strand of the genome, to generate the original pool of the potential sgRNA binding sites.
2. To avoid potential off-target effects, we mapped all 5’-XX XXXXX XXXXX-NGG-3’ from step #1 back to the genome using the short reads mapping program Seqmap (http://www-personal.umich.edu/~jianghui/seqmap/) with parameter setting “1 /output_all_matches”, and filtered out the sequences with multiple mappings.
3. We required that the designed sgRNAs should be able to fold properly. To check that this was true, we linked the 42nt scaffold sequence 5’-GTTTTAGAGCTAGAAATAGCAAGTTAAAATAAGGCTAGTCCG-3’ to the 3’ end of the 20 nt specific target binding sequence and checked the folding structure of this 62nt sequence by RNA secondary structure prediction using RNAfold (http://www.tbi.univie.ac.at/~ronny/RNA/) with default parameters. We only kept the ones that the scaffold region could fold to the hair-pin structure as reported in Jinek et al *Science* 2012.
4. Finally, we filtered out any sgRNA sequences containing the BsaI restriction site (GGTCTC), which we used for cloning purposes.

The sequences that passed these four steps composed our pool of potential sgRNAs. Next, we chose sgRNAs from the sgRNA pool to target all (1) annotated genes, (2) promoters and (3) TFBSs, according to RegulonDB.

1. sgRNAs that target coding sequences: We tried to collect 4 sgRNAs for each annotated gene in the *E. coli* genome. We implemented a recursive approach to select sgRNAs as close to the ATG as possible and on the non-template strand for each gene. We first looked at the first 50%. Next, we looked at the annotated 5’ UTR regions, and the ones close to the start codon where selected with higher priority. Finally, we looked at the last half of the CDS sequence, and chose the sites closer to the start codon with higher priority. By using this approach, 4281 genes could be targeted with 4 sgRNAs, 193 additional genes could be targeted by 1-4 sgRNAs, and 158 genes could not be targeted by any sgRNA. We further looked at the 158 genes that could not be targeted by the previous pipeline. We noticed that 39 of them were located in operons where an upstream gene in that operon had properly selected sgRNAs. For the rest 109 genes, we found many of them had closely related homologs on the genome, which caused the sgRNAs targeting these region to be not unique on the genome and could target both of the homologs. So we compared the sequences of all the annotated genes, and defined a homolog gene set by performing a megablast search with parameter setting of “-F F - D 3 -e 1e-10”. We searched for the potential sgRNA target sites that locate in both the homolog genes but not any other sites on the genome. 48 genes could be targeted in this way. Finally, there are 71 genes could not be targeted by our sgRNA design procedure. Most of them are small RNAs that don’t have any PAM site. Finally, we designed 17622 sgRNAs, which could target 4561 genes (4522 directly, 39 indirectly) on the *E.coli* genome.
2. sgRNA target promoters: For the promoters that did not overlap with any annotated UTR or CDS regions, we selected the sgRNA from both the sense and anti-sense strand in the region from upstream 60 bp to downstream 10 bp relative to the transcription start site. For the promoters located within a gene body, we only designed sgRNAs that binds to the template strand of that region. 14257 sgRNAs were selected to target 7404 Promoters.
3. sgRNA target TFBS sites: We designed all the sgRNAs that could target the TFBSs annotated in the RegulonDB database. An sgRNA is selected if it could cover at least one-third of the annotated TFBS. If the TFBS is shorter than 15 bp, we required that the overlap should be at least 5bp. 1867 sgRNAs were selected to target 1264 TFBS sites.
4. sgRNA for subcategories:

a. We designed sgRNAs for 21 genes subcategories (eg cell division, small RNAs, central intermediary metabolism). These sgRNAs are encoded with an additional category code in the 3’ end of each library oligo to enable amplification of subpools of the library. Categories and their corresponding category codes for amplification can be found in Supplementary Table 1.

We used the following external files as annotations for our sgRNA design:

- Genome sequence: Escherichia coli str. K-12 substr. MG1655, complete genome, NCBI Reference Sequence NC_000913.2 (http://www.ncbi.nlm.nih.gov/nuccore/NC_000913.2)
- Genome annotations from RegulonDB v8.1: (http://regulondb.ccg.unam.mx/download/Data_Sets.jsp)

∘ Gene coordinate: Gene_sequence.txt
∘ Promoter annotation: PromoterSet.txt
∘ UTR annotation: UTR_5_3_sequence.txt
∘ Transcription factor binding sites: BindingSiteSet.txt

Note: To keep with genome annotation updates, sgRNAs were remapped to promoter and TFBS features using more recent RegulonDB annotations:

- Promoter annotation: PromoterSet.txt (RegulonDB v9.4; release date 05-08-2017)
- TFBS annotation: BindingSiteSet.txt (RegulonDB v10.5; release date 09-13-2018)

## Supplementary Note 2. Analysis of time-series data

The fitness of each sgRNA strain was calculated at each sequenced time point relative to the initial timepoint of the experiment. This constructed a time-series fitness curve for each sgRNA in the library.

Time-series Analysis 1 – Clustering of Essential Genes:

1. Calculate gene fitness scores for each gene annotated as essential in the PEC database
2. Filter out any genes that did not have a gene fitness score ≤ -1 (i.e. keep only essential genes that showed a knockdown phenotype)
3. Keep only timepoints with a Pearson correlation ≥ 0.8 across two replicates
4. Average the remaining timepoints across replicates
5. Performed a min-max scaling of each timepoint (i.e. i.e. fitness values at each timepoint were scaled to between 0 and 1) from Step 4 to ensure that all timepoints were treated equally
6. Used the Elbow method to track the variation of the within-cluster-sum-of-squares (WCSS) with the number of clusters (k – ranging from 1 to 14) and found k = 3 to be the optimal number of clusters for K-means based on visual inspection.
7. Performed K-means clustering with selected k from Step 6 to classify essential gene curves
8. Visualize K-means clusters (Early / Mid / Late)

Time-series Analysis 2 – Clustering of All Genes:

1. Calculate gene fitness scores for each gene targeted in the CRISPRi library
2. Keep only timepoints with a Pearson correlation ≥ 0.8 across two replicates
3. Average the remaining timepoints across replicates
4. Performed a min-max scaling of each timepoint (i.e. i.e. fitness values at each timepoint were scaled to between 0 and 1) from Step 3 to ensure that all timepoints were treated equally
5. Used the Elbow method to track the variation of the within-cluster-sum-of-squares (WCSS) with the number of clusters (k – ranging from 1 to 14) and found k = 3 to be the optimal number of clusters for K-means based on visual inspection.
6. Performed K-means clustering with selected k from Step 5 to classify essential gene curves
7. Visualize K-means clusters (Early / Late / No Effect)

Gene Ontology Enrichment for Analysis 1 and 2:

For either time-series analysis, each gene was associated with its annotated TIGR Role. A hypergeometric test was carried out for each TIGR Role in each gene class (for analysis 1 – Early / Mid / Late; for analysis 2 – Early / Late / No Effect) with parameters: N = #total essential genes in data set, K = #total genes in class, n = #total genes with TIGR Role in data set, k = #genes with TIGR Role in class. The Benjamini-Hochberg correction was applied to the resulting p-values using the multitest function (parameter: “fdr_bh”) in the statsmodels python module (http://www.statsmodels.org/stable/index.html). The threshold of p_FDR-adjusted_ ≤ 0.05 was used as the significance threshold.

Time-series Analysis 3 – Comparison of gene-targeting and promoter-targeting CRISPRi:

1. Select all essential genes for which guides targeting the corresponding promoter and the gene itself were designed in the library
2. Of these promoter-gene pairs, select all essential genes that are the first and only gene in their respective transcription unit (TU). This enables association of a specific promoter knockdown or gene phenotype to the specific gene itself.
3. Of the remaining promoter-gene pairs, select cases where the gene only has one promoter
4. Keep only sgRNAs that had t0 counts ≥ 10 and had a fitness score ≤ -1 by the final timepoint (i.e. timepoint 15)
5. Plot time-series using lineplot function from seaborn plotting library (v0.9.0) with the parameter setting “ci = 95” to generate 95% confidence intervals via bootstrapping.
  a. Lineplot function: https://seaborn.pydata.org/generated/seaborn.lineplot.html
6. For each gene, compare the overlap of the 95% confidence intervals between population doublings 6 and 12 (these timepoints were selected because they are both highly correlated across replicates and because after doubling 12 we start to see fitness scores leveling out due to limitations in sequencing read depth)

